# Antiviral reverse transcriptases reveal the evolutionary origin of telomerase

**DOI:** 10.1101/2025.10.16.682844

**Authors:** Stephen Tang, Josephine L. Ramirez, Mario Rodríguez Mestre, Dennis J. Zhang, Megan Wang, Tanner Wiegand, Yanzhe Ma, Marko Jovanovic, Rafael Pinilla-Redondo, Samuel H. Sternberg

## Abstract

Defense-associated reverse transcriptases (DRTs) employ diverse and distinctive mechanisms of cDNA synthesis to protect bacteria against viral infection. However, much of DRT family diversity remains unstudied. Here we identify a new antiviral defense system, DRT10, that associates with a non-coding RNA (ncRNA) to catalyze processive, protein-primed synthesis of tandem-repeat DNA. Repeat addition is dictated by sequence and structural features of the ncRNA that have direct parallels in the RNA component of telomerase. Remarkably, a phylogenetic analysis of RTs across domains of life reveals an unexpected link between DRT10 and telomerase that is further supported by structural comparisons and mechanistic evidence. These findings expand the repertoire of reverse transcription mechanisms in antiviral defense and point to a bacterial origin for telomerase.

**One-Sentence Summary:** Insights from antiviral defense systems reveal an unexpected bacterial origin for the mechanism of chromosome maintenance by eukaryotic telomerase.

## INTRODUCTION

Reverse transcriptases (RTs) are abundant enzymes found across all domains of life (*1*). Their hallmark activity is the synthesis of complementary DNA (cDNA) from an RNA template, first discovered in retroviruses through the pioneering work of Baltimore and Temin (*2, 3*). In the decades since, dozens of additional RT families have been discovered, with diverse functions across an expansive range of biological contexts.

Although RTs likely originated in mobile genetic elements (*4*), they have been repeatedly domesticated and repurposed for specialized cellular functions (*5*). In eukaryotes, the best-studied example is the telomerase reverse transcriptase (TERT), which extends telomeric repeats at chromosome ends to mitigate replication-associated DNA loss (*6, 7*). In bacteria, most domesticated RTs characterized to date function in antiviral immunity (*8*). Our recent work on a family of systems termed defense-associated RTs (DRTs) uncovered strikingly diverse and unusual mechanisms by which these enzymes protect their hosts from bacteriophage infection (*9, 10*).

DRTs belong to a large, diverse collection of bacterial RTs termed the Unknown Group (UG) (*11*), which can be subdivided into three classes based on their phylogenetic relationships and associated protein domains (*12*). The first UG enzymes to be experimentally studied were the abortive infection proteins AbiK, AbiA, and AbiP2 (*13–15*), which belong to Class 1. Biochemical and structural studies showed that Abi RTs catalyze protein-primed, untemplated DNA synthesis (*16, 17*), though the functional roles of these DNA products in phage defense have not been elucidated. Class 2 UG enzymes, by contrast, carry out unique modes of RNA-templated DNA synthesis, as exemplified by DRT2 and DRT9 systems. DRT2 exploits a looped noncoding RNA (ncRNA) template for rolling circle reverse transcription to create an immune effector gene *de novo* (*9, 18*), while DRT9 repetitively reverse transcribes a poly-uridine tract within its ncRNA to yield toxic deoxyadenylate homopolymers (*10, 19*). Despite their vastly different outcomes, both systems employ iterative reverse transcription of a ncRNA template to generate long, repetitive cDNA species.

These findings suggest that additional Class 2 UG systems might also rely on ncRNA templates, and that repetitive reverse transcription to generate functional cDNAs could represent a unifying mechanistic feature. One candidate system, UG17, attracted our attention due to its frequent localization within defense islands (*12*), strongly suggesting a role in phage defense. We therefore set out to investigate the biological function and molecular mechanism of UG17. Here we show that UG17, hereafter named DRT10, associates with a ncRNA and SLATT domain-containing protein to provide robust defense against diverse phages. Reverse transcription is primed by a serine residue within the RT itself, and is templated by a well-defined region within the ncRNA to generate tandem-repeat cDNAs. Sequence and structural features of the ncRNA dictate repeat synthesis in a manner strikingly reminiscent of telomerase repeat addition. Remarkably, phylogenetic analyses reveal a close evolutionary link between DRT10 and TERT itself, suggesting that telomerase originated in bacteria before being co-opted for the maintenance of linear genomes in eukaryotes.

## RESULTS

### DRT10 is a tripartite phage defense system

UG17 RTs are encoded adjacent to SLATT domain-containing proteins and show evidence of tight co-evolution (**Fig. 1A and fig. S1A**). As UG17 RTs and SLATT proteins have both been documented to associate with known antiphage immune systems (*12, 20*), we hypothesized that UG17 RTs would also function in phage defense. We selected three distinct *E. coli* homologs of UG17, along with their associated SLATT proteins and native flanking sequences, to test for defense activity in the laboratory strain, *E. coli* MG1655 (**Fig. 1A and fig. S1B**). All three systems exhibited robust defense activity across a diverse panel of *E. coli* phages, as assessed by plaque assay (**Fig. 1B**). Defense was abolished with either deletion of the *SLATT* gene or mutation of the RT catalytic residues (**Fig. 1C and fig. S1C**), but deletion of a tyrosine recombinase associated with two of the three systems had no observable effect on immune activity (**fig. S1C**). These results establish that UG17 operons confer RT-dependent immunity against phage infection, and therefore we term this phage defense system DRT10, in line with established nomenclature (*11, 12*).

**Figure 1.**
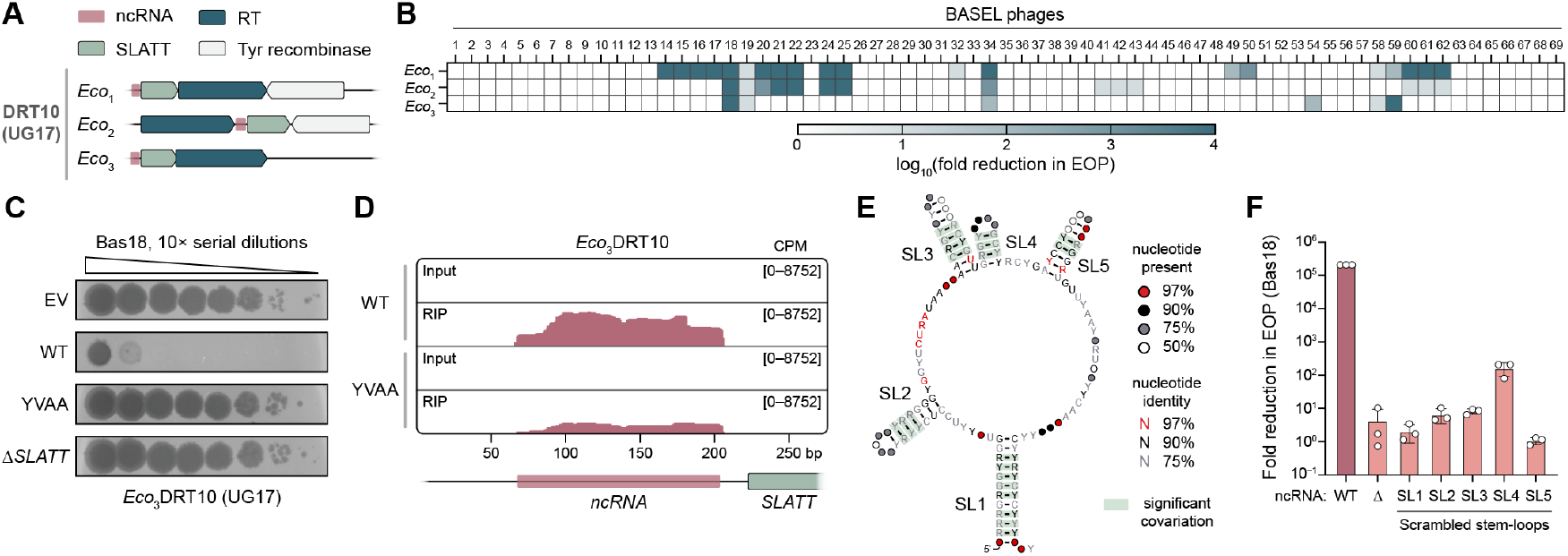
DRT10 is a tripartite phage defense system. **(A)** Schematic of representative DRT10 (UG17) operons. **(B)** Heatmap of phage defense activity for select DRT10 (UG17) operons tested against the BASEL collection of *E. coli* phages (*47*) quantified as the fold reduction in efficiency of plating (EOP) relative to an empty vector (EV) control. **(C)** Plaque assays demonstrating that *Eco*_3_DRT10 (*Eco*_3_UG17) defense activity against Bas18 phage is lost with mutation of the RT catalytic residues (YVDD>YVAA) or deletion of the *SLATT* gene (Δ*SLATT*). **(D)** RIP-seq coverage tracks for WT or catalytically inactive (YVAA) RT compared to non-immunoprecipitated (“Input”) controls, revealing a noncoding RNA (ncRNA) within the intergenic region upstream of SLATT. A schematic of the genetic locus from the *Eco*_3_DRT10 expression plasmid is shown below; data are normalized for sequencing depth and plotted as counts per million reads (CPM). **(E)** Covariance model of the *Eco*_3_DRT10 ncRNA secondary structure from an analysis of 307 homologous systems; conserved stem-loops (SL) are numbered. **(F)** Bar graph of *Eco*_3_DRT10 defense activity against Bas18 phage for the indicated ncRNA mutants, quantified as in **B**. Data are from *n* = 3 technical replicates.

We and others previously showed that DRT2 and DRT9 systems encode ncRNAs that harbor templates for reverse transcription (*9, 10, 18, 19*). We hypothesized that DRT10 systems would similarly encode RT-associated ncRNAs and therefore tested the *Eco*_3_DRT10 RT for RNA binding in cells. RNA immunoprecipitation and sequencing (RIP-seq) experiments with a FLAG-tagged RT showed strong enrichment of reads within an intergenic region upstream of the *SLATT* gene, for both the WT RT and a catalytically inactive mutant (**Fig. 1D)**, revealing the presence of a DRT10-associated, 137-nucleotide (nt) ncRNA. Using the experimentally determined boundaries of the *Eco*_3_DRT10 ncRNA, together with a collection of 307 DRT10 homologs, we next built a covariance model (CM) to evaluate the ncRNA for conserved sequence and structural features. This analysis yielded a consensus structure prediction that featured a large, central stem-loop (SL) decorated with four smaller stem-loops (**Fig. 1E**). In parallel, we performed RIP-seq experiments with *Eco*_1_DRT10 and *Eco*_2_DRT10, which similarly showed signal enrichment upstream of the *SLATT* gene and revealed ncRNAs with comparable secondary structures (**fig. S1D**,**E**). Notably, the predicted structures of the DRT10 ncRNAs closely resemble the known structure of the *Sen*DRT9 ncRNA (**fig. S1F**), highlighting their relatedness and suggesting similar functions in templating reverse transcription.

We proceeded to test the function of the DRT10 ncRNA in phage defense with a series of genetic perturbations. Deletion of the entire ncRNA predictably led to a complete loss of defense against Bas18 phage (**Fig. 1F**). To more subtly disrupt the ncRNA secondary structure without changing its overall nucleotide composition, we individually scrambled each predicted stem-loop. The resulting mutants also severely diminished defense activity, underscoring the importance of these ncRNA structural elements for DRT10 function (**Fig. 1F**).

Collectively, these results establish a biological function for UG17 reverse transcriptases in DRT10-mediated phage defense. DRT10 systems exhibit a tripartite architecture comprising a ncRNA, SLATT protein, and RT, all of which are required for antiviral protection.

### Protein-primed synthesis of repetitive cDNA

We next sought to discover DRT10 reverse transcription products. We developed a simple, scalable method, termed miniprep-seq, to enrich non-chromosomal DNA from cells for high-throughput sequencing (HTS) (**Materials and Methods**). This method recovered the major features of the DRT2 and DRT9 cDNAs previously revealed by cDNA immunoprecipitation and sequencing (cDIP-seq) (*9, 10*), including DRT2 rolling circle junctions and DRT9 homopolymer motifs (**fig. S2A**,**B**), establishing miniprep-seq as a facile and high-throughput alternative to the comparatively laborious cDIP-seq workflow. When we applied miniprep-seq to cells expressing *Eco*_3_DRT10 and aligned reads to the expression plasmid, we did not observe any clear signal over the ncRNA locus for the WT RT compared to the catalytically inactive control, initially suggesting that the ncRNA was not templating reverse transcription (**fig. S2C**). However, given the similarity of the DRT9 and DRT10 ncRNAs (**fig. S1F**), as well as our previous experience identifying DRT9 ncRNA-templated cDNAs by analyzing *unmapped* sequencing reads (*10*), we reasoned that a similar approach might again yield insights into unexpected reverse transcription products.

We proceeded to extract unmapped reads from *Eco*_3_DRT10 miniprep-seq datasets and perform motif discovery (**Fig. 2A**), searching for sequence patterns differentially enriched in the WT relative to the RT-inactive condition. Strikingly, this analysis revealed a statistically significant, 9-nt motif (5ʹ–GAAT-CATTG–3ʹ) that was present in 5.9% of unmapped reads for the WT condition, compared to only 0.13% for the mutant RT (**Fig. 2B**). Inspection of the ncRNA immediately revealed a 9-nt segment within the central loop region that bore the reverse complement sequence (**Fig. 2C**), implicating this region as the likely cDNA template. Although identification of the 9-nt DNA motif and its ncRNA reverse complement suggested that our earlier read mapping should have revealed cDNA signal, we reasoned that this seemingly contradictory result might be explained if the motif was embedded within longer sequences that failed to match the corresponding upstream and downstream regions of the template. To test this possibility, we extracted all instances of the 9-nt motif within unmapped reads and built a consensus sequence logo of their neighboring context. Remarkably, this analysis revealed that the 9-nt motif was flanked by direct repeats of the motif itself (**Fig. 2D**). Reads containing multiple repeats of the motif were exclusively found in the WT, but not the mutant, RT condition, whereas reads containing only a single repeat — which would arise simply from plasmid sequencing — were equally abundant in both conditions (**fig. S2D**). Further analyses revealed that the majority of unmapped reads in the WT RT condition contained at least 15 consecutive repeats of the 9-nt motif (**Fig. 2E**), just below the maximum limit of 16 repeats for a 150-cycle sequencing read.

**Figure 2.**
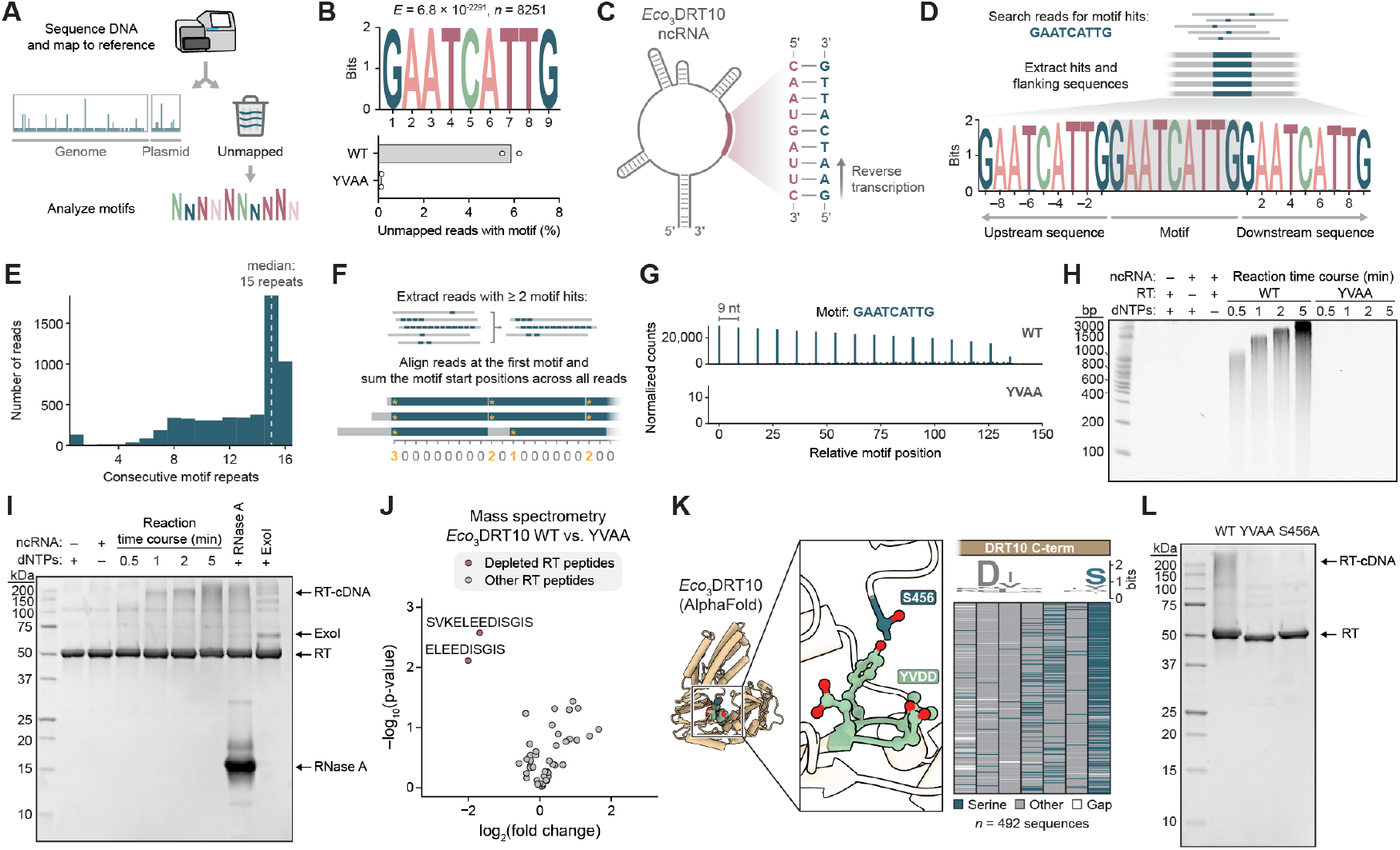
Protein-primed synthesis of tandem-repeat cDNA by DRT10. **(A)** Schematic of unmapped read analysis workflow. **(B)** *Top*: MEME motif discovery results for unmapped miniprep-seq reads from cells expressing *Eco*_3_DRT10, showing the top enriched motif for WT samples relative to a catalytically inactive (YVAA) control. *E* represents the *E*-value significance; *n* represents the number of contributing sites. *Bottom*: Percentage of unmapped reads that contain the 9-nt motif. Data are shown for *n* = 2 independent biological replicates. **(C)** Illustration of the *Eco*_3_DRT10 ncRNA secondary structure; the inset highlights the putative template region that harbors the reverse complement of the DNA motif identified in **B. (D)** *Top*: Schematic of workflow to extract motif hits and analyze their sequence context. *Bottom*: Sequence logo of motif hits and their flanking neighborhood from *Eco*_3_DRT10 miniprep-seq data, showing that motif hits are present as direct repeats. **(E)** Histogram of motif-containing reads from an *Eco*_3_DRT10 miniprep-seq dataset, quantifying the longest continuous stretch of motif repeats per read. The distribution median is 15 repeats of the 9-nt motif; 16 repeats is the maximum possible value for a 150-nt sequencing read. **(F)** Schematic of method to visualize the position and quantity of motif repeats in sequencing reads. **(G)** Motif position graph for miniprep-seq datasets from *Eco*_3_DRT10 WT and YVAA samples, generated as shown in **F**. Data are normalized as counts per million plasmid-mapping reads for each sample. **(H)** Denaturing 15% urea-PAGE analysis of *in vitro* RT activity assays that investigated the minimal components and time dependence of DRT10 cDNA synthesis. Reactions contained 2.5 µM RT, 2.5 µM ncRNA, and 100 µM dNTPs, and were incubated at 37 °C for 5 minutes unless otherwise indicated. Reactions were terminated by boiling and treated with proteinase and RNase prior to electrophoretic analysis; sizes from a dsDNA ladder are marked. **(I)** Denaturing 12% SDS-PAGE analysis of *in vitro* RT activity assays. Reactions contained 2.5 µM RT, 2.5 µM ncRNA, and 100 µM dNTPs, and were incubated at 37 °C for 5 minutes unless otherwise indicated. Sizes from a protein ladder are marked. **(J)** Volcano plot of differential peptide abundance from mass spectrometry analysis of immunoprecipitated *Eco*_3_DRT10 RT proteins (WT vs. YVAA). Each dot represents an RT peptide arising from tryptic cleavage, and red dots indicate significantly depleted peptides with log_2_(fold change) ≤ –2 and p-value < 0.05, as determined by an unpaired two-tailed t-test. The two depleted peptides are distinct tryptic cleavage products that both derive from the RT C-terminus. **(K)** AlphaFold 3 structure prediction of the *Eco*_3_DRT10 RT (left), with an inset (middle) showing the close proximity of the C-terminal serine (S456) to the RT catalytic residues (YVDD). A multiple sequence alignment of 492 DRT10 RT homologs (right) highlights the strong conservation of a serine residue at this position. **(L)** Denaturing 12% SDS-PAGE analysis of *in vitro* RT activity assays with WT RT or the indicated mutants, performed as in **I**. Sizes from a protein ladder are marked.

To better visualize the repeat pattern and abundance of cDNA motifs, we searched all unmapped reads for the 9-nt motif and recorded the start coordinate of each hit, relative to the first hit within the same read. We then summed up the number of hits at every position across the 150-nt read length within a given sample and plotted the results as a “motif position graph” (**Fig. 2F**). A precise 9-nt periodicity was clearly evident in the motif position graph for *Eco*_3_DRT10 (**Fig. 2G**), highlighting the tandem-repeat nature of the cDNA products. Miniprep-seq analysis of *Eco*_1_DRT10 and *Eco*_2_DRT10 systems revealed similar tandem-repeat motifs templated from the analogous region of their respective ncRNAs, with notable variability in the motif sequence, motif length, and precision of repeat addition (**fig. S2E**,**F**).

Next, we assessed how phage infection affects DRT10-mediated cDNA synthesis. *Eco*_3_DRT10 produced the same tandem-repeat cDNA in Bas18-infected cells as in uninfected cells, but at reduced levels (**fig. S2G**). This result contrasts with DRT2 and DRT9 immune systems, both of which exhibit increased cDNA production upon phage infection (*9, 10, 18, 19*). Since phage infection triggers second-strand cDNA synthesis in DRT2 systems (*9*), we also searched for evidence of complementary strand synthesis by DRT10; however, second-strand cDNA products were barely detectable in both uninfected and infected conditions (**fig. S2H**). We conclude that *Eco*_3_DRT10 constitutively generates single-stranded DNA (ssDNA) products whose abundance diminishes during phage infection, though whether this decrease is integral to the defense mechanism or a by-product of infection remains unclear.

To define the minimal molecular requirements for cDNA synthesis, we biochemically reconstituted RT activity *in vitro* from individual components. After purifying the *Eco*_3_DRT10 RT (**fig. S3A**) and incubating it with dNTPs and chemically synthesized ncRNA, we performed denaturing urea-PAGE analysis of the reaction products, which revealed a time-dependent, high-molecular-weight smear that formed only when the RT, ncRNA, and dNTPs were present (**Fig. 2H**). DNA sequencing confirmed that *in vitro* products exhibited tandem repeats of the 9-nt motif, matching *in vivo* miniprep-seq results (**fig. S3B**), and mutations to the RT active site abolished activity, as expected (**Fig. 2H**). DNA synthesis was also sensitive to dNTP concentration and availability of Mg^2+^ ions (**fig. S3C**), further establishing these kilobase-length species as *bona fide* reverse transcription products of *Eco*_3_DRT10.

We next probed the chemical nature of the cDNA product by subjecting it to various enzymatic treatments. The cDNA was efficiently degraded by a ssDNA-specific endonuclease (Nuclease P1) but not by a dsDNA-specific endonuclease (Duplex DNase) (**fig. S3D**), consistent with *Eco*_3_DRT10 synthesizing ssDNA. Interestingly, exonuclease treatment yielded different results depending on enzyme polarity: a 3ʹ-5ʹ exonuclease (ExoI) completely degraded the product, whereas a 5ʹ-3ʹ exonuclease (RecJ_f_) had no effect (**fig. S3D**). This led us to hypothesize that the 5ʹ end was protected by an RNA or protein primer, as has been reported for several other bacterial RTs (*10, 18, 21, 22*). Omission of the usual RNase A pre-treatment step prior to electrophoretic separation had no effect on product mobility, whereas omission of proteinase K treatment prevented the DNA product from efficiently entering the gel (**fig. S3E**), strongly suggesting the presence of a covalent protein–DNA linkage. This was further supported by SDS-PAGE analysis of the RT reaction products, which revealed the emergence of a high-molecular-weight smear upon dNTP incubation that was unaffected by RNase treatment but largely eliminated by DNase treatment (**Fig. 2I**).

To identify candidate RT residues involved in protein-primed reverse transcription, we immuno-precipitated the WT or catalytically inactive RT from *E. coli* cells and analyzed peptides by mass spectrometry, hypothesizing that a covalent DNA linkage would interfere with detection of peptides containing the priming residue, as shown previously for DRT9 (*10*). Strikingly, the only peptides that were significantly depleted in WT relative to RT-mutant samples mapped to the RT C-terminus (**Fig. 2J**). We identified a highly conserved serine residue (S456) within this region that was predicted by AlphaFold 3 to be positioned near catalytic residues within the active site (**Fig. 2K**), suggesting that its hydroxyl side chain could serve as the priming nucleophile. In agreement with this model, mutation of the serine to alanine (S456A) severely impaired cDNA synthesis *in vitro* and *in vivo* and completely abolished phage defense activity (**Fig. 2L and fig. S3E-G**). We obtained similar results in biochemical reactions with *Eco*_1_DRT10 (**fig. S3H-J**), for which the likely C-terminal priming residue is a tyrosine (Y444), establishing protein-primed DNA synthesis as a shared feature among diverse DRT10 RTs. All other examples of protein-primed reverse transcription use tyrosine residues (*10, 17, 21, 23, 24*), to our knowledge, further highlighting *Eco*_3_DRT10 as the first example of a serine-primed reverse transcriptase.

### RNA determinants of tandem-repeat cDNA synthesis

Having established that the DRT10 RT and ncRNA are necessary and sufficient for protein-primed reverse transcription, we next sought to determine how this system efficiently generates tandem-repeat cDNAs with high precision. Guided by our previous work on DRT9 — where single-nt substitutions across the ncRNA loop containing the template (hereafter “template loop”) largely abolished phage defense (*10*) — we hypothesized that the sequence context around the DRT10 template might similarly influence repeat addition, and in turn affect phage defense. We therefore systematically mutated the *Eco*_3_DRT10 template loop by substituting each position with its complementary nucleotide, and then tested the mutants for defense activity. Unexpectedly, the ncRNA template loop was highly tolerant of mutagenesis, with only five of twenty substitutions eliminating defense (**Fig. 3A**).

**Figure 3.**
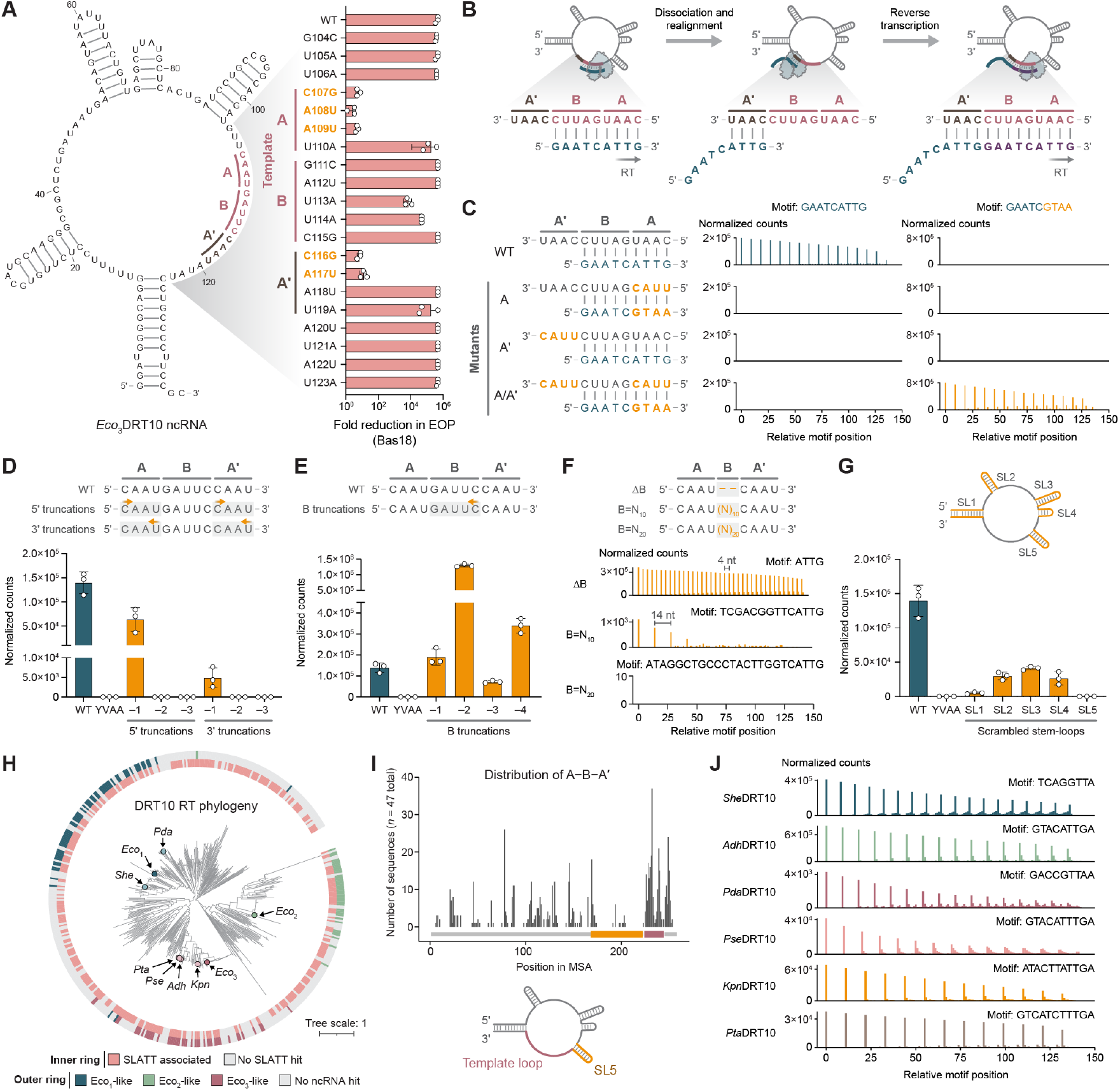
A ncRNA repeat pattern programs tandem-repeat cDNA synthesis. **(A)** *Left*: Predicted secondary structure of the *Eco*_3_DRT10 ncRNA, with coordinates numbered based on the 137-nt species identified by RIP-seq. An A–B–Aʹ repeat pattern is highlighted, in which a variable region (B) is flanked by repeated sequences (A and Aʹ). The template (bolded red) corresponds to A–B, while the downstream repeat (bolded brown) corresponds to Aʹ. *Right*: Bar graph of defense activity against Bas18 phage for the indicated *Eco*_3_DRT10 template loop substitution mutants, quantified as the fold reduction in EOP relative to an EV control. Defense-inactivating mutations are bolded in orange. Data are mean ± SD for *n* = 3 technical replicates. **(B)** Mechanism of *Eco*_3_DRT10 repeat addition. Reverse transcription of the template terminates at the 5ʹ template boundary, and the nascent cDNA then dissociates from the template and reanneals at Aʹ, providing a primer for the subsequent round of reverse transcription. Each cycle extends the cDNA by a 9-nt repeat corresponding to the reverse complement of A–B. **(C)** *Left*: Schematic of ncRNA template region mutants that disrupt and rescue A–Aʹ homology. *Middle*: Motif position graphs from mini-prep-seq datasets of the mutants schematized at left, quantifying the cDNA repeat motif templated by the WT ncRNA. *Right*: Motif position graphs quantifying the cDNA repeat motif templated by the mutant ncRNAs. Data are normalized as counts per million plasmid-mapping reads for each sample. **(D)** *Top*: Schematic of ncRNA template region mutants that incrementally truncate A and Aʹ from either the 5ʹ or 3ʹ end. *Bottom*: Bar graph quantifying the number of miniprep-seq reads containing ≥ 3 consecutive cDNA motif repeats for the mutants schematized at top, alongside WT and catalytically inactive (YVAA) controls. Data are normalized as in **C**, and represent the mean ± SD for *n* = 3 independent biological replicates. **(E)** *Top*: Schematic of ncRNA template region mutants that incrementally truncate B from the 3ʹ end. *Bottom*: Bar graph quantifying miniprep-seq reads as in **D. (F)** *Top*: Schematic of ncRNA template region mutants that vary the length of B, including complete deletion of B (ΔB), replacement of B with a synthetic 10-mer (B=N_10_), and replacement of B with a synthetic 20-mer (B=N_20_). *Bottom*: Motif position graphs quantifying the indicated cDNA repeat motifs in miniprep-seq reads, normalized as in **C. (G)** *Top*: Cartoon of the *Eco*_3_DRT10 ncRNA secondary structure highlighting stem-loop (SL) structures that were disrupted by mutations. *Bottom*: Bar graph quantifying miniprep-seq reads for the indicated SL mutants as in **D**, alongside WT and YVAA controls. **(H)** Phylogenetic tree of clustered DRT10 RT representatives, showing SLATT associations (inner ring) and ncRNA hits (outer ring). Branches corresponding to experimentally tested candidates are highlighted. **(I)** Distribution of bioinformatically predicted A–B–Aʹ motifs in *Eco*_3_-like ncRNA sequences. Motif midpoints are plotted non-redundantly, visualizing one motif midpoint per ncRNA. **(J)** Motif position graphs from miniprep-seq datasets of the indicated DRT10 homologs, quantifying the cDNA repeat motifs predicted by A–B–Aʹ identification. Data are normalized as in **C**.

The observation that six positions within the putative cDNA template could be altered without loss of defense suggested that DRT10 immunity can tolerate alternative DNA product sequences. Indeed, miniprep-seq of the U110A mutant revealed robust synthesis of a tandem-repeat product precisely matching the altered template (**fig. S4A**). Conversely, the observation that two positions outside of the putative template were critical for defense activity suggested that they were important to the mechanism of cDNA synthesis. Miniprep-seq confirmed that both mutations (C116G and A117U) severely diminished tandem-repeat synthesis, as did the three defense-inactivating mutations within the template (**fig. S4A**). Comparison of the inactive mutants drew our attention to an intriguing feature of the template and its downstream sequence: the first four bases of the template exactly matched the four bases immediately downstream of the template. We denote this sequence pattern as A–B–Aʹ, where A–B is the template and Aʹ is the downstream segment identical to A. Notably, all five mutations associated with loss of defense and cDNA repeat synthesis created mismatches between A and Aʹ (**Fig. 3A and fig. S4A**).

Taken together, these observations allowed us to postulate the following mechanism for DRT10-mediated tandem-repeat cDNA synthesis: Protein-primed reverse transcription initiates using A–B as a template, and is followed by dissociation of the nascent cDNA from the ncRNA template. Re-annealing of the cDNA at Aʹ then provides a DNA primer for a subsequent round of reverse transcription, and iterative cycles of extension, dissociation, and template resetting append additional 9-nt repeats (**Fig. 3B**). Remarkably, our mutational data precisely aligned with this model: mutating the first two bases of A or Aʹ abolished repeat addition, whereas mutating the fourth base was tolerated (**Fig. 3A and fig. S4A**), reflecting a more stringent base-pairing requirement at the 3ʹ end of the DNA primer. Mutation of the third base produced asymmetric effects: the Aʹ mutation (which would cause a T:U mismatch) was tolerated, whereas the A mutation (which would cause an A:A mismatch) was not (**Fig. 3A and fig. S4A**), consistent with the greater steric clash from a purine–purine mismatch. Closer inspection of the sequencing results from mutations to the fourth base of A and Aʹ revealed another intriguing asymmetry: the mutation in A (U110A) yielded a cDNA with a mutated repeat sequence, while the mutation in Aʹ (U119A) preserved the WT repeat sequence (**fig. S4A**). This result demonstrates that templating of cDNA synthesis is dictated by the A–B segment, whereas realignment of the cDNA during repeat addition is mediated by the Aʹ segment.

We comprehensively tested the model by systematically perturbing the A–B–Aʹ pattern and assessing repeat addition by miniprep-seq. Mutating A or Aʹ alone completely eliminated repeat addition, whereas mutating them in combination restored repeat addition with a correspondingly altered repeat sequence (**Fig. 3C**), demonstrating that A–Aʹ homology is necessary and sufficient for repeat synthesis. To define the minimum length requirement for A and Aʹ, we simultaneously truncated both features in single-nt increments and found that loss of a single nucleotide from the 5ʹ or 3ʹ end of A and Aʹ was tolerated, but further truncations were not (**Fig. 3D**). These results indicate that *Eco*_3_DRT10 generally requires at least three bases of A–Aʹ homology for efficient template resetting, though the results from our single-nt substitution panel suggest that two bases of homology are in some cases sufficient (see above).

In contrast to A and Aʹ, we anticipated that sequence requirements for B would be more flexible. Indeed, all tested truncation variants — including complete removal of B — were tolerated, with some variants even exceeding WT levels of cDNA synthesis activity (**Fig. 3E,F**). Conversely, increasing the length of B revealed an upper limit: low-level repeat addition persisted when substituting the native 5-mer with a synthetic 10-mer, but was undetectable with substitution of a 20-mer (**Fig. 3F**). Finally, we assessed the contributions of larger structural elements in the ncRNA by applying miniprep-seq to the scrambled stem-loop (SL) variants previously tested for phage defense. All SL perturbations decreased repeat addition, but only mutation of SL5 completely eliminated repeat products (**Fig. 3G**). Given that SL5 lies immediately upstream of the template loop, these data suggest a role for SL5 in setting the 5ʹ template boundary.

We next asked whether this repeat addition templating mechanism generalizes to homologous DRT10 immune systems. Intriguingly, our analysis of DRT10-encoded ncRNAs revealed poor sequence conservation within the template loop (**Fig. 1E and fig. S1E**), unlike the universally conserved poly-uridine template region in DRT9 systems (*10*). Nonetheless, focusing first on *Eco*_1_DRT10 and *Eco*_2_DRT10, we were able to identify A–B–Aʹ patterns within both template loops that matched their respective cDNA repeat products (**fig. S4B**, compare with **fig. S2E**,**F**). The A and Aʹ sequences are only 2-nt long for both ncRNAs, indicating that A–Aʹ homology lengths can vary between systems. Furthermore, inspection of the A–B–Aʹ sequence for *Eco*_1_DRT10, which exhibits less precise repeat addition than the other homologs (**figs. S2E, S4C**,**D**), revealed a cryptic re-annealing site within the B segment that likely interferes with accurate resetting of the template (**fig. S4E**). In support of this, a mutation to the B segment that removed this site dramatically improved repeat periodicity (**fig. S4F**).

Our proposed repeat addition templating mechanism would predict that diverse DRT10-associated ncRNAs should preserve characteristic A–B–Aʹ patterns, in spite of considerable primary sequence variation. We therefore assembled a dataset of diverse ncRNAs (**Fig. 3H, Tables S1**,**S2**), and aligned the resulting CM hits. Our analysis revealed that A–B–Aʹ patterns were highly enriched within the template loop (**Fig. 3I, fig. S4G**,**H, Table S3**), with a substantial fraction (91.5%) of *Eco*_3_DRT10-like ncRNAs harboring an identifiable A–B–Aʹ pattern within this region, compared to 95.0% and 85.0% of *Eco*_1_DRT10-like and *Eco*_2_DRT10-like ncRNAs, respectively. Finally, to experimentally validate A–B–Aʹ predictions, we selected six additional DRT10 systems for gene synthesis and heterologous testing in *E. coli*. Strikingly, analysis of cDNA production by miniprep-seq revealed tandem-repeat cDNAs for all six systems that exactly matched our A–B–Aʹ predictions, across a wide range of motif lengths and sequence compositions (**Fig. 3J**).

We conclude that DRT10 repeat synthesis relies on defined sequence and structural features of the associated ncRNA. While DRT10 systems naturally accommodate a wide range of template sequences, two constraints appear to be inflexible: (1) an A–B–Aʹ repeat pattern that facilitates template realignment and (2) positioning of the template downstream of a stable stem-loop.

### An evolutionary link to repetitive DNA synthesis in eukaryotes

As we deciphered the mechanism of cDNA synthesis by DRT10 immune systems, we noticed striking parallels to the mechanism of repeat addition by eukaryotic telomerase enzymes (**Fig. 4A**) (*25*). Linear eukaryotic chromosomes shorten with each round of replication, leading to genome instability and cellular senescence (*26*). Telomerase mitigates this progressive loss of DNA — known as the “end-replication problem” — by extending the repetitive telomeric sequences that cap chromosome termini (*27*).

**Figure 4.**
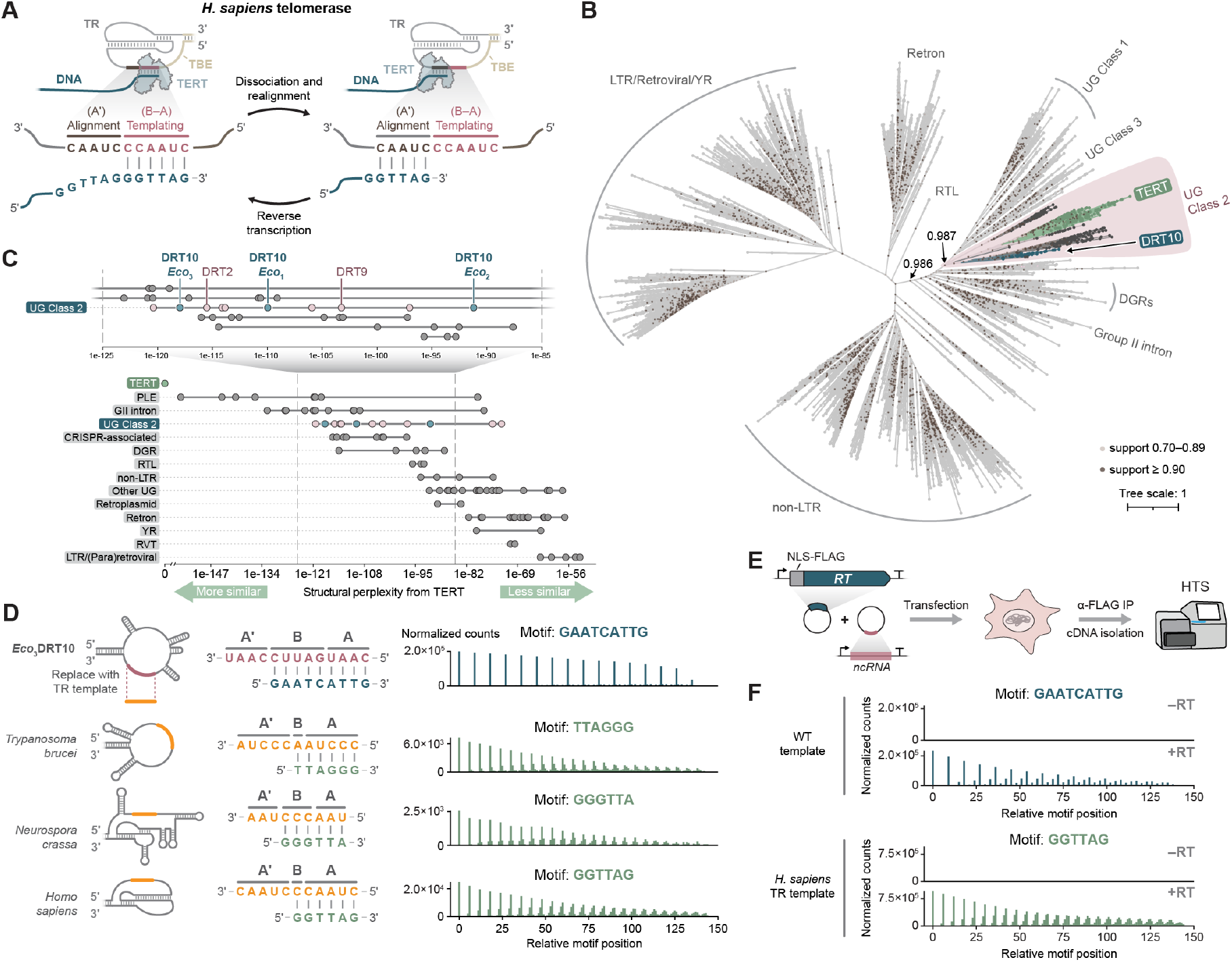
An evolutionary link between DRT10 and telomerase. **(A)** Illustration of the *H. sapiens* telomerase catalytic cycle that extends chromosome ends. The core RT (TERT, light blue) associates with a telomerase RNA (TR, gray) to add cDNA repeats to telomeres by repetitively reverse transcribing a short template region (red) within the TR. Following extension through the 5ʹ end of the templating segment, which is defined by a structured template boundary element (TBE, brown), TERT and the nascent cDNA dissociate and reanneal at the alignment segment to prime a subsequent cycle of cDNA synthesis. The templating segment is analogous to A–B within the DRT10 ncRNA, and the alignment segment is analogous to Aʹ. **(B)** Phylogenetic tree of major RT families across the three domains of life. The core RT domain (motifs 1–7) from 6,734 representative sequences was used to build the unrooted tree (**Table S4, Data S1-S3**). RTs broadly cluster into LTR, non-LTR, and bacterial subfamilies. TERT is located in a well-supported clade within the bacterial cluster (local support value: 0.987), together with UG RTs including DRT10. **(C)** Structural perplexity analysis assessing structural similarity between selected RT proteins (**Tables S5**,**S6**) and human TERT. Each dot represents the perplexity value of the comparison between the selected structure retrieved from the AlphaFold Database (or when unavailable, predicted by AlphaFold 3) and human TERT. Lower perplexity values indicate greater similarity. **(D)** *Left*: Schematic of template substitution experiments in which the A–B–Aʹ region of the *Eco*_3_DRT10 ncRNA was replaced with the analogous region from the telomerase RNA of *T. brucei, N. crassa*, or *H. sapiens*. The expected cDNA repeat products, corresponding to the known telomeric DNA repeat sequences for each organism, are shown. *Right*: Motif position graphs from miniprep-seq datasets of *E. coli* cells expressing *Eco*_3_DRT10 with the chimeric ncRNAs, compared to a WT control (top). Data are normalized as counts per million plasmid-mapping reads for each sample. **(E)** Schematic of cDIP-seq experiments assessing reconstitution of DRT10-mediated cDNA repeat addition in human cells. **(F)** Motif position graphs from cDIP-seq datasets of HEK293T cells expressing *Eco*_3_DRT10 with either the WT ncRNA template or the *H. sapiens* TR template, plotted as in **D**. Data are normalized for sequencing depth as counts per million reads, and no-RT control conditions are shown.

The telomerase ribonucleoprotein complex comprises an RT (TERT) and a separately encoded telomerase RNA (TR). Despite substantial divergence among TRs, two features are broadly conserved: (1) a short template region composed of distinct “alignment” and “templating” segments, and (2) a template boundary element (TBE) formed by a stem-loop or pseudoknot that helps define the template 5ʹ end (*28*). Both features are critical in enabling repeat addition processivity (RAP), the hallmark feature of telomerase enzymatic activity, whereby multiple telomeric repeats (i.e., cDNA motifs) are synthesized within a single RT–substrate binding event (*29, 30*). During each cycle, reverse transcription of the TR templating segment is followed by dissociation and re-annealing of the nascent cDNA to the alignment segment, thereby priming the next round of cDNA synthesis (**Fig. 4A**). Intriguingly, the key RNA features that govern telomerase RAP have direct parallels in the DRT10 ncRNA: the templating and alignment segments correspond to the A–B and Aʹ sequences, respectively, and the TBE corresponds to the essential SL5 element upstream of A–B–Aʹ.

The profound mechanistic overlap between DRT10 and telomerase led us to wonder whether the similarity might reflect shared ancestry rather than convergent evolution. We therefore set out to investigate the evolutionary relationship between TERT and DRT10 RTs, as well as their broader placement within RT phylogeny. We assembled a representative set of 6,734 RTs spanning all experimentally characterized families and performed a phylogenetic analysis of their RT domains (**Fig. 4B, Table S4, Data S1-S3**). The resulting tree recovered established phylogenetic groupings, with the major eukaryotic (non-LTR and LTR retrotransposon) and prokaryotic (retron, group II intron, and DGR) RTs all forming monophyletic clades. A separate, well-supported clade (local support value: 0.987) encompassed most UG RTs and, remarkably, all 181 TERT homologs in our dataset (**Fig. 4B**). Penelope-like element (PLE) RTs, which constitute another family of eukaryotic retroelements, also localized to this clade (**fig. S5**), consistent with their previously documented relatedness to TERT (*31*). The placement of TERT and PLEs alongside UG RTs was especially striking given the clear partitioning of other eukaryotic families into their own separate clades. Importantly, the overall topology of the tree was recapitulated by orthogonal, structure-based comparisons (**fig. S6, Data S4**).

Closer inspection of the tree revealed that, within the clade encompassing TERT, PLE, and UG RTs, the members were divided into two major subclades (**Fig. 4B and fig. S5**). One subclade primarily comprised Class 1 and Class 3 UG RTs, while the other primarily comprised Class 2 UG RTs, including DRT10, alongside TERT and PLE RTs (**Fig. 4B and fig. S5**). This suggested that TERT might be more closely related to Class 2 UG RTs than to Class 1 and Class 3. However, because relationships between highly diverged proteins can be difficult to resolve from primary sequence alone, we complemented the phylogeny with an information-theoretic measure of structural similarity (*32*). We retrieved structure predictions for 160 representative RTs from the AlphaFold Database and quantified their structural perplexity to human TERT (hTERT; **Tables S5**,**S6**). Among UG RTs, the nine lowest perplexity scores (indicating greater structural similarity) belonged to Class 2 enzymes, including DRT2, DRT9, and DRT10 (**Fig. 4C**), supporting a close relationship between TERT and Class 2 UG RTs. Interestingly, group II intron RTs also exhibited low perplexity, suggesting similarities to TERT that were less evident at the primary sequence level. Among all RTs included in this analysis, PLE RTs were most similar to hTERT, whereas non-LTR and LTR retrotransposon RTs were more distant, in agreement with the results from phylogenetic analysis.

Finally, considering the mechanistic similarities and phylogenetic relatedness between DRT10 and telomerase, we sought to test their functional compatibility. We replaced the *Eco*_3_DRT10 ncRNA template sequence with TR templates from *Trypanosoma brucei, Neurospora crassa*, and *Homo sapiens* — each of which directs the synthesis of a uniquely permuted, canonical telomeric TTAGGG repeat — and assessed cDNA synthesis in *E. coli*. Remarkably, all three chimeric ncRNA species generated abundant cDNA products matching the telomeric sequences from each organism (**Fig. 4D**). These results furthermore extended to human cells, where co-transfection of expression vectors for the *Eco*_3_DRT10 RT and ncRNA containing the *H. sapiens* TR template elicited robust synthesis of human telomeric DNA repeats (**Fig. 4E-F and fig. S7**). These results underscore the functional similarity of DRT10 and telomerase repeat addition mechanisms, while also highlighting the flexibility of DRT10 as a platform for repetitive DNA synthesis in diverse cellular contexts.

Collectively, our findings point to the ancient origin of a telomerase-like repeat addition mechanism in bacteria that likely predated the emergence of linear chromosomes. In prokaryotes, this mechanism was further elaborated to protect cells against genetic parasites, while in early eukaryotes, it was co-opted to protect linear genomes against replication-associated DNA loss (**Fig. 5**).

**Figure 5.**
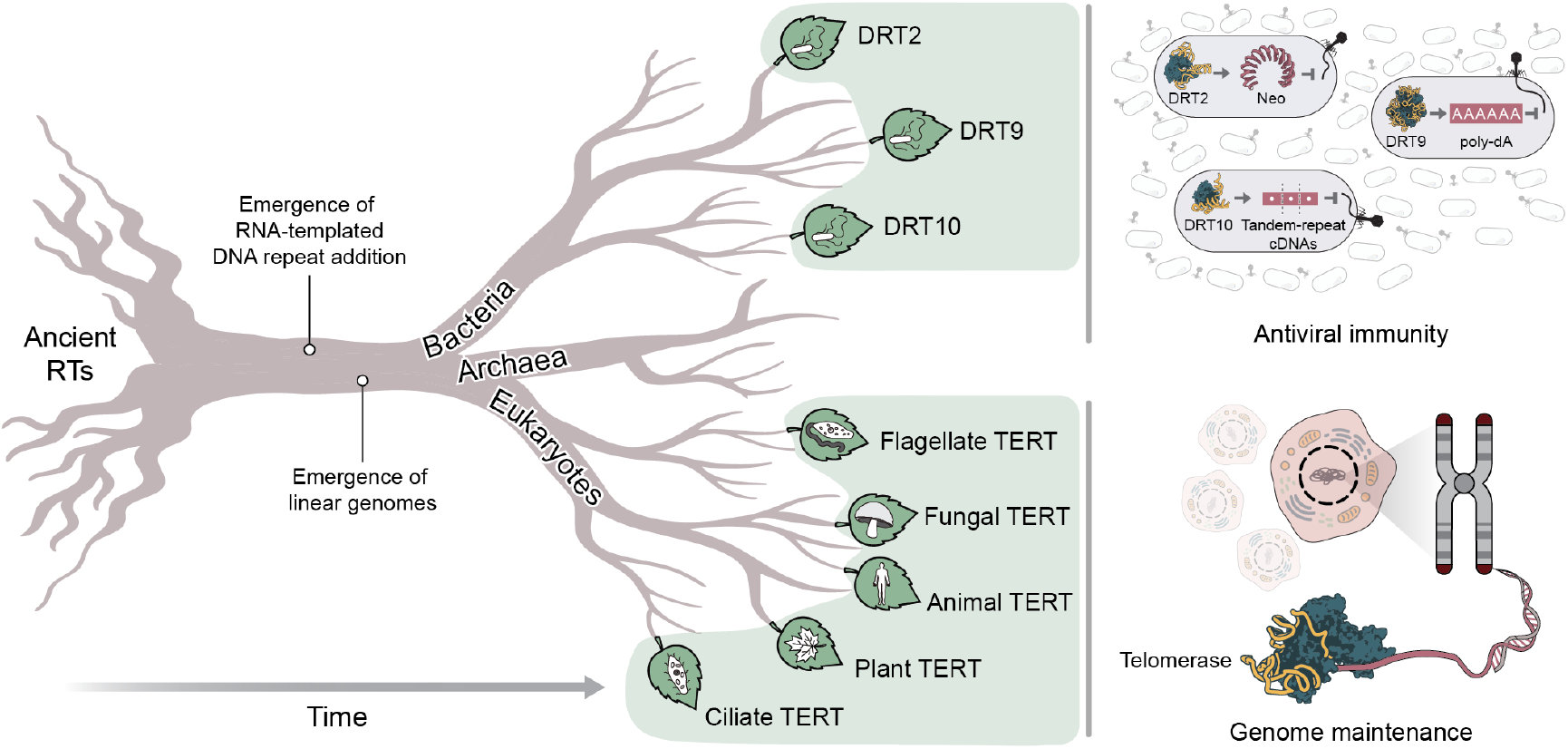
Evolutionary origins and diversification of UG RTs and telomerase. Proposed evolutionary trajectory of UG RTs, including those encoded by DRT2, DRT9, and DRT10, and telomerase RT (TERT) homologs across the eukaryotic domain. RNA-templated DNA repeat addition first arose in an ancient bacterial RT, and was later acquired and co-opted by eukaryotes as a fundamental process for maintaining linear genomes. In prokaryotes, telomerase-like repeat addition diversified into multiple RT-based strategies for antiviral immunity.

## DISCUSSION

Our discovery of RNA-templated cDNA repeat addition in DRT10 immune systems provides a direct functional and evolutionary link between antiviral immunity in bacteria and a fundamental genome maintenance mechanism in eukarya. Phage defense is conferred through a distinctive mechanism of DNA repeat synthesis catalyzed by the DRT10-encoded RT and its associated ncRNA, and further requires the presence of a SLATT protein. Reverse transcription is primed from a serine residue within the RT itself, creating a covalent protein–DNA linkage, and proceeds through iterative cycles of dissociation and realignment directed by a hallmark A–B–Aʹ sequence pattern in the ncRNA template. The repeat addition mechanism of DRT10 closely parallels that of telomerase reverse transcriptase (TERT), and phylogenetic analyses reveal an unexpected evolutionary relationship, suggesting that ancient TERT-like enzymes arose in bacteria and persist today in a diverse family of antiviral reverse transcriptases.

These findings shed new light on the long-debated evolutionary origin of telomerase (*33*). Earlier models suggested that telomerase might have evolved in eukaryotes through the domestication of a mobile retroelement RT, such as those encoded by non-LTR retrotransposons (*34*). However, telomerase differs from these enzymes in key aspects: while retrotransposon RTs reverse transcribe their own full-length mRNAs in *cis*, telomerase repetitively reverse transcribes a short template embedded within a separately encoded ncRNA gene. Previous phylogenetic analyses of RTs across domains of life have yielded inconsistent topologies with respect to TERT (*34–37*), though these analyses notably did not include DRT10 or other UG RT sequences. Incorporating these sequences now resolves the placement of TERT with high confidence, which is further supported by structural comparisons and mechanistic evidence. Our results indicate that TERT is more closely related to bacterial RTs than to other major eukaryotic RT clades, though how TERT was ultimately exapted by eukaryotes remains intriguingly unresolved. One possibility is horizontal gene transfer via a mobile ancestor shared with PLE RTs, its closest known eukaryotic relatives (*31*); alternatively, an ancestral, non-mobile telomerase could have been acquired by endosymbiosis prior to the emergence of eukarya. Discriminating between these potential scenarios will require further study.

The cellular function of the ancient bacterial progenitor to modern telomerase remains unknown — and may never be fully resolved. Although some bacteria possess linear chromosomes, these are typically capped by “protelomerase” enzymes that generate covalently closed DNA hairpins but bear no ancestral relationship to telomerase (*38*). But given that phage–host conflict has persisted for billions of years, it is plausible that the telomerase precursor would have also functioned in antiviral defense. All experimentally characterized UG enzymes to date have been implicated in phage defense (*12*), pointing to a conserved antiviral role for this enzyme family and its ancestral members.

Within this apparent functional uniformity lies a striking diversity of mechanisms by which RNA-mediated, repetitive DNA synthesis has been elaborated into distinct strategies of phage defense. Repetitive reverse transcription by DRT2 assembles a promoter upstream of a start codon to elicit expression of a cryptic immune gene (*9, 18*), while the analogous process in DRT9 drives the synthesis of toxic, kilobase-length DNA homopolymers (*10, 19*). DRT10 generates tandem-repeat cDNA products whose immune role remains unclear, but likely involves co-encoded SLATT effector proteins. Although the RNA templates and DNA products of most UG systems remain uncharacterized, our findings suggest that telomerase-like cDNA repeat addition may represent a unifying theme across all or part of the UG RT family, adapted in diverse ways for phage defense.

Despite their commonalities and close evolutionary relationships, UG RTs and telomerase exhibit a surprising diversity of molecular architectures. DRT2 functions as a monomer and primes cDNA synthesis from the 3ʹ end of its ncRNA (*18*), whereas DRT9 assembles into a hexamer and primes from internal tyrosine residues, covalently linking the cDNA product to the RT (*10*). TERT, by contrast, acts as a monomer and primes off of substrate DNA, and relies on several domains in addition to its core RT domain for interactions with its DNA substrate and RNA template (*39, 40*). DRT10 closely resembles TERT in its repeat synthesis mechanism, but it primes from a C-terminal serine residue and is considerably more compact (*Eco*_3_DRT10 is 456 amino acids whereas hTERT is 1,132 amino acids), suggesting that it may represent a streamlined, minimal ortholog of TERT. Further work will be needed to provide structural insights into DRT10-mediated antiviral immunity, and to resolve the molecular details of additional RT-based defense systems within UG Classes 1–3.

A key question that remains unanswered by our study is how the tandem-repeat cDNA product of DRT10 contributes to immune function. An emerging theme across bacterial immune systems is the regulation of membrane-associated effectors by nucleic acid-based signaling molecules (*41, 42*). Within the DRT family, several other systems — including DRT1, DRT3, and DRT9 — have been noted to co-occur with predicted membrane proteins (*11, 12*). Given that SLATT proteins are predicted to form membrane pores (*43*), and that DRT10 RT and SLATT proteins exhibit tight co-evolution (**Fig. S1A**), we propose that the DRT10 cDNA product directly modulates SLATT activity, potentially as a signaling molecule to control a toxic membrane effector. The repetitive nature of the cDNA may enable multivalent engagement of an oligomeric SLATT complex, with each repeat unit bound by an individual SLATT monomer. Although the defensive function of such an assembly remains speculative, our finding that phage infection reduces DRT10 cDNA abundance raises the possibility that SLATT toxicity is neutralized by the cDNA in uninfected cells and then unleashed to drive abortive infection in the presence of phage.

Telomerase is found in nearly all eukaryotes, where it carries out an essential role in chromosome maintenance. Loss of telomerase in somatic cells is a hallmark of aging (*44*), while aberrant telomerase reactivation underlies many cancers (*45*). By linking telomerase to DRT-family enzymes, our work reveals its evolutionary origin while also opening new avenues for exploring its functions. Studies of other bacterial immune systems have revealed direct counterparts in human immunity and shown how bacterial pathways can illuminate poorly understood aspects of their human analogs (*46*), and the same may hold true for DRT systems and telomerase. Beyond these biological insights, our study also demonstrates that DRT10 can be reprogrammed to synthesize tandem-repeat cDNAs of custom sequence, including human telomeric DNA. This activity suggests that DRT10 could in principle be harnessed to explore new strategies addressing telomere dysfunction in aging and disease.

## ACKNOWLEDGMENTS

We thank A.I. Palmieri, A.J. Robinson, and T.M. Smith for laboratory support, A. Harms for generously providing the BASEL phage collection, and the JP Sulzberger Columbia Genome Center for NGS support.

## AUTHOR CONTRIBUTIONS

S.T. and S.H.S. conceived the project. S.T. performed most experiments and experimental analyses. J.L.R. performed phage defense assays, RIP-seq experiments, and biochemical reconstitution of DRT10-mediated tandem-repeat cDNA synthesis. M.R.M. assembled and curated reverse transcriptase datasets and performed RT phylogenetic analyses, in close collaboration with D.J.Z. and with supervision from R.P.-R. M.W. helped with cloning and performed plaque assays for ncRNA mutants. T.W. performed DRT10 phylogenetic analyses, analyzed ncRNAs for A–B–A′ patterns, performed analyses of protein priming, and assisted with data visualization. Y.M. performed mass spectrometry experiments to identify the protein priming residue, with supervision from M.J. S.T. and S.H.S. discussed the data and wrote the manuscript, with input from all authors.

## DATA AND MATERIALS AVAILABILITY

Next-generation sequencing data are made available in the National Center for Biotechnology Information (NCBI) Sequence Read Archive. Datasets generated and analyzed in the current study are available from the corresponding author upon reasonable request.

## SUPPLEMENTARY MATERIALS

Materials and Methods

Supplementary Text

Figures S1 to S7

Tables S1 to S9

Data S1 to S4

