## Supplementary Materials for "Antiviral reverse transcriptases reveal the evolutionary origin of telomerase"

**The PDF file includes:**

Materials and Methods

Supplementary Text

Figs. S1 to S7

**Other Supplementary Materials for this manuscript include the following:**

Tables S1 to S9

Data S1 to S4

### MATERIALS AND METHODS

Plasmid and *E. coli* strain construction

All strains and plasmids used in this study are described in **Tables S7 and S8**. DRT10 operons, including their native upstream and downstream flanking sequences, were chemically synthesized (GenScript) and cloned into a pACYC184 plasmid backbone by Gibson Assembly. Derivative plasmids were cloned using a variety of methods, including around-the-horn PCR, restriction digestion-ligation, and Golden Gate Assembly. All plasmids were cloned and propagated in *E. coli* strain NEB Turbo (sSL0410) and verified by Sanger sequencing or whole plasmid sequencing. All DRT10 homologs deriving from taxa other than *Enterobacteriaceae* were cloned downstream of a synthetic promoter and ribosome binding site (RBS) to ensure strong heterologous expression in *E. coli*. Due to overlap between the *SLATT* and *RT* genes in the native $Eco_3$DRT10 operon, a stuffer sequence and synthetic RBS were added upstream of the *RT* in the N-terminally FLAG-tagged RT construct used for immunoprecipitation experiments. Miniprep-seq experiments in **Figs. 3C-G**,**4D** were performed using $Eco_3$DRT10 expression plasmids lacking the *SLATT* gene. For ncRNA expression in human cells, mutations were introduced to disrupt poly-U tracts that would partially terminate transcription by RNA polymerase III (**fig. S7**).

Manual inspection of the multiple sequence alignment for closely related DRT10 homologs revealed a likely misannotation in the translation start site for $Eco_3$DRT10 RT (NCBI accession OKX54664), leading to the inclusion of two extra amino acids at the N-terminus. Numbering of the amino acid sequence in the manuscript has been modified to reflect this (i.e., S458 in the NCBI record is numbered here as S456).

Phage propagation and plaque assays

Bacteriophages were propagated in liquid culture by picking single plaques and inoculating LB media, supplemented with 5 mM $CaCl_2$ and 5 mM $MgSO_4$, containing *E. coli* MG1655 cells diluted 1:100 from overnight cultures. After 4–5 hours of incubation with shaking at 37 °C, lysates were centrifuged at 4,000 x g for 10 min to pellet cell debris, and chloroform was added to a final concentration of 5% to lyse residual bacteria. Supernatants were passed through a sterile 0.22 μm filter and stored at 4 °C.

Small-drop plaque assays were performed as previously described (*9*). Briefly, *E. coli* K-12 strain MG1655 (sSL0810) was transformed with the indicated plasmids, and individual colonies were used to inoculate LB media and were grown overnight at 37 °C. Cultures were mixed with molten soft agar (0.5% agar in LB media supplemented with 5 mM $CaCl_2$ and 5 mM $MgSO_4$) at 45 °C and poured over solid bottom agar (1.5% agar in LB media containing the appropriate antibiotic) in a Petri dish. 10× serial dilutions of phage in LB media were spotted onto the surface of the soft agar lawn, and plates were incubated at 37 °C for 8–16 hours to allow plaque formation. Plaque forming units (PFU) $mL^{-1}$ were calculated using the following formula: . When individual plaques were indistinguishable, a count of 50 plaques was assigned at the lowest dilution showing visible lawn clearance. Phage defense activity was assessed by calculating the fold reduction in efficiency of plating (EOP), which was determined by dividing the PFU $mL^{-1}$ obtained on a lawn of empty vector (EV)-transformed control cells by the PFU $mL^{-1}$ obtained on a lawn of defense system-expressing cells.

To screen DRT10 homologs against the BASEL phage collection (*47*), plaque assays were performed as described above, but with assistance from a Beckman Coulter Biomek FX liquid handler. Transformants of *E. coli* K-12 strain MG1655 were inoculated in liquid LB media containing the appropriate antibiotic and grown overnight at 37 °C with shaking. The next day, overnight cultures were mixed with molten soft agar at 45 °C and poured over solid bottom agar in a rectangular Petri dish. 10× serial dilutions of phage in LB media were made using the Biomek liquid handler 8-channel pipetting arm and spotted onto the surface of the soft agar lawn using the 96 head channel pipettor. Plates were incubated at 37 °C for 8–16 hours to allow plaque formation.

<u>RNA immunoprecipitation and sequencing (RIP-seq) from *E. coli* cells</u>

RIP-seq in *E. coli* was performed as previously described (*9*). *E. coli* K-12 strain MG1655 (sSL0810) cells were transformed with DRT10 plasmids encoding N-terminally 3xFLAG-tagged RT proteins with a WT or mutant (YVAA) active site. Individual colonies were used to inoculate 20 mL cultures in liquid LB media and were grown at 37 °C to mid-log phase. Cells were harvested by centrifugation at 4,000 × *g* for 10 min at 4 °C and then washed with 5 mL of cold TBS (20 mM Tris-HCl pH 7.5 at 25 °C, 150 mM NaCl), prior to being centrifuged again at 4,000 × *g* for 5 min at 4 °C. Pellets were then washed with 1 mL of cold TBS and centrifuged at 10,000 × *g* for 5 min at 4 °C. After removing supernatants, pellets were flash-frozen in liquid nitrogen and temporarily stored at –80 °C.

To prepare antibody–bead complexes for immunoprecipitation, Dynabeads Protein G (Thermo Fisher Scientific) were washed 3× in 1 mL IP Lysis Buffer (20 mM Tris-HCl pH 7.5 at 25 °C, 150 mM KCl, 1 mM $MgCl_2$, 0.2% Triton X-100), resuspended in 1 mL IP Lysis Buffer, combined with anti-FLAG antibody (Sigma-Aldrich, F3165), and rotated for > 3 hours at 4 °C. 30 µL of beads and 10 µL of antibody mix were prepared per sample. Antibody-bead complexes were washed 2× to remove unbound antibodies and resuspended in IP Lysis Buffer to a final volume of 30 µL per sample.

Flash-frozen pellets were thawed on ice and resuspended in 1.2 mL IP Lysis Buffer supplemented with 1× Complete Protease Inhibitor Cocktail (Roche) and 0.1 U $\mu L^{-1}$ SUPERase•In RNase Inhibitor (Thermo Fisher Scientific). Cells were lysed by sonication, and lysates were cleared by centrifugation at 21,000 × *g* for 15 min at 4 °C. Supernatants were transferred to new tubes, and a 10 µL aliquot of each sample was set aside and stored at –80 °C as an "input" control. The remainder of each sample was combined with 30 µL of antibody-bead complex and rotated overnight at 4 °C. The next day, each sample was washed 3× with 1 mL ice-cold IP Wash Buffer (20 mM Tris-HCl pH 7.5 at 25 °C, 150 mM KCl, 1 mM $MgCl_2$). For elution from beads, supernatants were removed, and beads were resuspended in 750 µL TRIzol (Thermo Fisher Scientific). After 5 min incubation at RT, supernatants containing eluted RNA were transferred to new tubes. Samples were then mixed with 150 µL chloroform, incubated at 25 °C for

2 min, and centrifuged at 12,000 × *g* for 15 min at 4 °C. RNA was isolated from the upper aqueous phase using the RNA Clean & Concentrator-5 kit (Zymo Research). RNA from input samples was isolated in the same manner using TRIzol and column purification.

To prepare libraries for high-throughput sequencing, RNA was diluted in NEBuffer 2 and heated at 92 °C for 2 min for fragmentation by alkaline hydrolysis. Samples were then treated with TURBO DNase (Thermo Fisher Scientific), RppH (NEB), and T4 PNK (NEB) to remove DNA and prepare RNA ends for adapter ligation. RNA was purified using the Zymo RNA Clean & Concentrator-5 kit. Adapter ligation, reverse transcription, and indexing PCR were performed using the NEBNext Small RNA Library Prep kit. Libraries were sequenced on an Element AVITI in single-end mode with 150 cycles.

Miniprep-seq from *E. coli* cells

*E. coli* K-12 strain MG1655 (sSL0810) cells were transformed with the indicated plasmids, plated on LB agar with the appropriate antibiotic, and grown at 37 °C. Individual colonies for each replicate experiment were used to inoculate cultures in liquid LB media and grown overnight at 37 °C. The next day, overnight cultures were diluted 1:100 in 4 mL liquid LB media with the appropriate antibiotic and grown at 37 °C to $OD_{600}$ ~ 0.4. For experiments +/– phage infection, parallel cultures were grown, and phage was added to the infected condition at a multiplicity of infection (MOI) of 5. For phage T5, infections were allowed to proceed for 45 minutes, whereas for phage Bas18, infections were allowed to proceed for 90 minutes. Cells were harvested by centrifugation at 4,000 × *g* for 5 min, and the supernatant was removed. Cell lysis and DNA extraction was performed following the standard procedure for the QIAprep Spin Miniprep Kit (QIAGEN), with elution in 30 µL nuclease-free water.

To prepare high-throughput sequencing libraries, samples were heated at 95 °C for 2 min and then immediately placed on ice to denature DNA, enabling ligation of sequencing adapters to linear, ssDNA fragments. Adapter ligation and conversion of ssDNA to dsDNA were performed using the xGen ssDNA & Low-Input DNA Library Prep Kit (IDT) with half the recommended reaction volumes. PCR was performed to add in-house indexed primers using the NEBNext Ultra II Q5 Master Mix. Libraries were sequenced on an Element AVITI in single-end mode with 150 cycles.

RIP-seq analysis

RIP-seq datasets were processed as previously described (*9*). Cutadapt (*48*) (v4.2) was used to remove adapter sequences, trim low-quality ends from reads, and filter out reads shorter than 15 bp. For *E. coli* samples, reads were mapped to combined reference files containing the MG1655 genome (NC_000913.3) and relevant plasmid sequence using bwa-mem2 (*49*) (v2.2.1), with default parameters. SAMtools (*50*) (v1.17) was used to sort and index alignments. Coverage tracks were generated using bamCoverage (*51*) (v3.5.1) with normalization for sequencing depth, based on the total number of reads passing initial trimming and length filtering. Coverage tracks were visualized in IGV (*52*).

Miniprep-seq analysis

Sequencing reads were stringently trimmed and filtered with cutadapt to remove all residual adapter sequences, as previously described (*10*), with error, overlap, quality, and length thresholds set at -e 0.2 -O 10 -q 30 -m 45. Trimmed and filtered reads were mapped to combined reference files containing the MG1655 genome (NC_000913.3), phage genome for infected samples (T5: NC_005859.1; Bas18: MZ501095), and relevant plasmid sequence using bwa-mem2 (v2.2.1), with default parameters. Aligned reads were sorted, indexed, and plotted onto coverage tracks, as described above for RIP-seq analysis.

DRT2 cDNA repeat junction-spanning reads were quantified as previously described (*9*) by aligning reads to a custom reference sequence consisting of two concatenated cDNA repeats and using featureCounts (*53*) (v2.0.2) to count reads assigned to an annotation overlapping the junction. Analysis of unmapped reads in DRT9 miniprep-seq datasets was performed as previously described (*10*). Unmapped reads extracted from BAM alignment files using SAMtools fasta were analyzed using MEME (*54*) (v5.5.7) in differential enrichment mode with -minw 25 -maxw 150 -mod anr, using unmapped reads from miniprep-seq of the catalytically inactive RT (YAAA) as the control sequence set. Quantification of homopolymer-containing reads was performed as described (*10*), using countPattern from the Biostrings package (v2.70.3) in R to quantify reads containing at least 25 consecutive A, T, G, or C bases, allowing 1 mismatch. Reads containing homopolymers were counted and normalized according to the total number of filtered reads for each sample.

Unmapped reads in $Eco_3$DRT10 miniprep-seq datasets were extracted using SAMtools fasta. For MEME motif discovery and analysis of sequences flanking motif hits, unmapped reads were randomly downsampled to 10,000 reads per sample using seqtk v1.5-r133. MEME analysis was performed in differential enrichment mode with -minw 9 -maxw 30 -mod anr, using unmapped reads from miniprep-seq of the catalytically inactive RT (YVAA) as the control sequence set. Miniprep-seq datasets for $Eco_3$DRT10 and $Eco_2$DRT10 were processed similarly, with -minw 8 and -minw 5 for MEME analysis, respectively. Sequences flanking motif hits and the length distribution of tandem-repeat cDNAs were analyzed using custom scripts in R. Briefly, after motif hits were identified using matchPattern from the Biostrings package, the sequence of the *n* bases upstream and downstream of those coordinates — where *n* is the length of the motif — was recorded. Sequence logos were generated with ggseqlogo (*55*) v0.2 using the 'bits' method. To assess tandem-repeat cDNA length, sequencing reads were searched for consecutive motif hits using matchPattern. For each read, the longest run of uninterrupted motif repeats was determined. The distribution of repeat lengths was then summarized across all reads in a sample by plotting a histogram with ggplot2.

"Motif position graphs" were generated using custom scripts in R. Briefly, reads were scanned for exact matches to a specified DNA motif using matchPattern (Biostrings). To reduce background from plasmid-derived sequences, only reads containing at least 2 motif hits were retained. For each qualifying read, the start position of every motif hit was recorded relative to the first motif hit within that read. Relative motif positions were then aggregated across all reads in a sample, and counts of reads containing a motif at each relative position were tabulated. Motif counts were normalized for sequencing depth by scaling to the number of plasmid-mapping reads for each sample (obtained using SAMtools idxstats).

To quantify the abundance of reads with tandem-repeat cDNAs, reads were scanned for the presence of three consecutive repeats of the specified motif — chosen to reduce background from spurious matches, particularly for shorter motifs — using vcountPattern (Biostrings) with a Hamming distance of 1. Reads containing at least one such match were counted for each sample, and the resulting counts were normalized for sequencing depth by scaling to the number of plasmid-mapping reads for that sample.

To evaluate the precision of cDNA repeat addition for different DRT10 homologs, we calculated a "direct repeat fraction." For each sample, reads were searched for exact matches to the specified motif using matchPattern. Motif hits were classified as falling within a direct repeat if they were immediately preceded and followed by another copy of the exact same motif. The direct repeat fraction was calculated as the proportion of motif hits that were directly flanked on both sides, relative to the total number of motif hits. Hits that were too close to either end of the read to be identified as part of a direct repeat were excluded from the analysis.

DRT10 RT purification

To purify the RT from $Eco_3$DRT10 and $Eco_1$DRT10 systems, we cloned $His_6$-GST-tagged RT expression constructs (WT or mutant) and transformed *E. coli* BL21-AI cells. After growing cultures in LB media at 37 °C to $OD_{600}$ ~0.6, protein over-expression was induced with 0.25 mM IPTG, and cells were allowed to continue growing overnight at 16 °C. Cells were pelleted by centrifugation at 3,000 × *g* for 25 min at 4 °C, prior to lysis by sonication in Lysis Buffer (20 mM HEPES-KOH pH 8.0, 500 mM NaCl, 5 mM $MgCl_2$, 5% glycerol, 10 mM imidazole, 0.1% Triton-X-100, 1 mM TCEP, 1X cOmplete EDTA free Protease Inhibitor cocktail). The lysate was clarified by centrifugation at 10,000 × *g* for 30 min at 4 °C. Clarified lysates were then incubated for one hour with Ni-NTA resin at 4 °C (1 mL of Ni-NTA bead volume per 1 L initial culture), before being loaded into an Econo-Pac column (BioRad) where resin was allowed to settle. The resin was rinsed with 10 column volumes of Wash Buffer (20 mM HEPES-KOH, pH 8.0, 500 mM NaCl, 5 mM MgCl2, 5% glycerol, 20 mM Imidazole, 1 mM TCEP) prior to elution with Elution Buffer (20 mM HEPES-KOH pH 8.0, 500 mM NaCl, 5 mM MgCl2, 5% glycerol, 300 mM imidazole, 0.1% Triton-X-100, 1 mM TCEP). Desired fractions were combined and subjected to TEV protease cleavage for 5 hours at 4 °C, using 2.5 μg TEV protease per 100 μg target protein. Immediately following TEV cleavage, samples were diluted to 300 mM NaCl before performing ion exchange chromatography using a HiTrap Heparin 5 mL column (Cytiva). Samples were eluted from the column using a continuous gradient from Buffer A (20 mM HEPES pH 7.5, 300 mM NaCl, 1 mM TCEP, 5% glycerol) to Buffer B (20 mM HEPES pH 7.5, 1 M NaCl, 1 mM TCEP, 5% glycerol). The desired fractions were combined and concentrated using a 10 kDa MWCO PES Spin-X UF concentrator (Corning), before snap freezing in liquid nitrogen for long term storage at −80 °C. Protein purity was assessed by denaturing 10% SDS-PAGE analysis.

Biochemical cDNA synthesis assays

Chemically synthesized ncRNA (IDT) was folded by heating to 95 °C and ramp cooling down to 25 °C at a rate of 5 °C per minute. Reverse transcription reactions were then assembled by combining 2.5 µM RT and 2.5 µM ncRNA in Polymerization Buffer (50 mM Tris-HCl, pH 8.0, 100 mM NaCl, 2 mM MgCl2, 5 mM TCEP) and incubating for 10 min at 37 °C to promote RT–ncRNA complex formation. cDNA synthesis was initiated by the addition of dNTPs, and reactions proceeded at 37 °C for 60 min unless otherwise indicated. Reactions were terminated by heating to 95 °C for 2 minutes, prior to treatment with various nuclease or proteinase reagents, as indicated. Samples treated with Proteinase K (Thermo Fisher Scientific) were incubated at 55 °C for 30 min. All other enzymatic treatments were carried out at 37 °C for 30 min. Samples were boiled at 95 °C for 2 min in between each post-reaction enzymatic treatment.

DNA products from biochemical cDNA synthesis assays were visualized by 15% denaturing urea-PAGE. Samples were mixed in equal volumes with 2× RNA Loading Dye (95% formamide, .025% Bromophenol blue, .025% SDS), denatured by heating at 95 °C for 2 min, and then loaded onto gels for electrophoretic separation. Gels were stained using SYBR Gold (Thermo Fisher Scientific).

Protein components of biochemical cDNA synthesis assays were visualized by 12% denaturing SDS-PAGE. Samples were mixed with 6× SDS Loading Dye (375 mM Tris-HCl, pH 7.0, 9% SDS, 50% Glycerol, 0.03% Bromophenol blue) prior to denaturation by heating and electrophoretic separation. Gels were stained with SimplyBlue SafeStain (Thermo Fisher Scientific).

Sequencing of biochemical cDNA synthesis reaction products

DNA from biochemical cDNA synthesis was isolated using the Monarch Spin PCR and DNA Cleanup kit (NEB), following the Oligonucleotide Cleanup protocol. Adapter ligation and conversion of ssDNA to dsDNA was performed using the xGen ssDNA & Low-Input DNA Library Prep Kit (IDT) with half the recommended reaction volumes. Libraries were sequenced on an Element AVITI in single-end mode with 150 cycles. The resulting sequencing dataset was randomly downsampled to 100,000 reads using seqtk, and a motif position graph was generated as described above for miniprep-seq samples.

Sequence identity matrices

Sequence identity between DRT10 RT and SLATT homologs was analyzed by performing MAFFT alignment of amino acid sequences in Geneious using default settings. Accession numbers for RT and SLATT proteins are listed in **Table S9**.

<u>Immunoprecipitation-mass spectrometry (IP-MS)</u>

*Immunoprecipitation:*

*E. coli* K-12 strain MG1655 cells were transformed with plasmids encoding either WT or catalytically inactive (YVAA) *Eco*$_3$DRT10 RT containing an N-terminal 3xFLAG tag. Three individual colonies per condition were used to inoculate liquid LB media, and cultures were grown at 37 °C with shaking. Overnight cultures were diluted 1:100 in 50 mL fresh LB media and grown to $OD_{600}$ of 0.3. Cultures were infected with Bas18 phage at an MOI of 5 and grown for an additional 45 min at 37 °C. Cells were harvested by centrifugation at 4,000 × *g* for 10 min at 4 °C, washed with 1 mL of cold TBS, and centrifuged again at 15,000 × *g* for 1 min at 4 °C. After discarding supernatants, pellets were flash-frozen in liquid nitrogen and temporarily stored at –80 °C.

Antibody–bead complexes were prepared for immunoprecipitation by washing Dynabeads Protein G (Thermo Fisher Scientific) three times in 1 mL IP-MS Lysis Buffer (50 mM Tris-HCl pH 7.5, 150 mM NaCl, 5% glycerol, 1% IGEPAL CA-630). Washed beads were combined with anti-FLAG antibody (Sigma-Aldrich, F3165) and rotated for > 3 hours at 4 °C. 100 µL of beads and 10 µL of antibody were prepared per sample. Antibody-bead complexes were washed twice to remove unbound antibodies and were resuspended in IP-MS Lysis Buffer to a final volume of 100 µL per sample.

Flash-frozen pellets were thawed on ice and resuspended in 1.2 mL IP-MS lysis buffer supplemented with 1× Complete Protease Inhibitor Cocktail (Roche) and 0.1 U $\mu L^{-1}$ SUPERase•In RNase Inhibitor (Thermo Fisher Scientific). Cells were lysed by sonication and cell debris was cleared by centrifugation at 21,000 × *g* for 15 min at 4 °C. Supernatants were transferred to new tubes, and protein concentrations were measured by BCA assay (Thermo Fisher Scientific). For each sample, 1 mg of protein was combined with 100 µL of antibody-bead complex and rotated overnight at 4 °C. The next day, each sample was washed twice with 1 mL cold IP-MS Wash Buffer 1 (50 mM Tris-HCl, pH 7.5, 150 mM NaCl, 5% glycerol, 0.05% IGEPAL-CA-630) followed by another 2 washes with cold IP-MS Wash Buffer 2 (50 mM Tris-HCl pH 7.5 at 25 °C, 150 mM NaCl, 5% glycerol). On-bead tryptic digestion was performed in 80 µL Tris-urea buffer (50 mM Tris-HCl pH 7.5, 2 M urea, 1 mM DTT) with 5 $\mu g\ mL^{-1}$ sequencing grade modified trypsin (Promega), and incubated 1 h at 25 °C with regular agitation. Supernatants were transferred to new tubes, and beads were watched twice with 60 µL Tris-urea buffer without trypsin. Supernatants were combined with those from the first elution step, resulting in a total eluate volume of 200 µL per sample. Eluates were centrifuged at 5,000 x g for 1 min to remove residual beads, and supernatants were transferred to new tubes. Samples were flash-frozen in liquid nitrogen and temporarily stored at –80 °C prior to further processing.

*MS sample preparation:*

The eluates of immunopurified samples were reduced with 5 mM dithiothreitol (DTT) for 45 min at 25 °C while shaking (600 rpm), followed by alkylation in the dark using 10 mM iodoacetamide (IAA) for 45 min at 25 °C while shaking (600 rpm). Samples were then digested using 0.5 µg of Promega sequencing grade modified trypsin (Promega: V5111) at 25 °C and 600 rpm overnight. Digested peptides were acidified using formic acid, desalted on in-house packed C18 StageTips (two plugs), dried down using a Thermo Savant SpeedVac, and reconstituted in 3% acetonitrile/0.2% formic acid.

*LC-MS/MS analysis on a Q-Exactive HF:*

Digested peptides from the immunopurified samples were analyzed on a Waters M-Class UPLC using a 15 cm x 75 µm IonOpticks C18 1.7 µm column coupled to a benchtop Thermo Fisher Scientific Orbitrap Q Exactive HF mass spectrometer. Peptides were separated at a 400 nL $min^{-1}$ flow rate with a 90-min gradient, including sample loading and column equilibration times, using Solvent A (0.1% formic acid in water) and Solvent B (0.1% formic acid in acetonitrile). The detailed gradients were: 2% B for 1 min; linear increase to 10% B over 29 min; linear increase to 22% B over 27 min; linear increase to 30% B over 5 min; linear increase to 60% B over 4 min; linear increase to 90% B over 1 min; hold for 2 min; linear decrease to 50% B over 1 min, hold for 5 min; linear decrease to 2% B over 1 min; and re-equilibration at 2% B for 14 min.

Data were acquired in data-dependent mode using Xcalibur software, and each cycle's 12 most intense peaks were selected for MS2 analysis. MS1 spectra were measured with a resolution of 120,000, an AGC target of 3e6, and a scan range from 300 to 1800 m/z. MS2 spectra were measured with a resolution of 15,000, an AGC target of 1e5, a scan range from 200–2000 m/z, and an isolation window width of 1.6 m/z.

Raw data were searched against a combined reference proteome that included the *E. coli* K-12 strain MG1655 proteome, Bas18 phage proteome, and the plasmid-encoded $\mathit{Eco}_3$DRT10 RT and SLATT sequences, using MaxQuant (*56*) (v2.0.3.0). MaxQuant was run with default settings, with the label minimum ratio count set to 1. Label-free quantification was enabled, and both iBAQ and "match between runs" features were turned on.

For peptide-level analysis to compare peptide detection between the immunoprecipitated WT and YVAA RT samples, data were first filtered to remove peptides with LFQ intensity of 0. Values were then normalized to the sum of LFQ intensities for each condition. To identify significantly depleted peptides in the WT compared to the YVAA condition, the $\log_2$(fold change) was calculated along with p-values by unpaired two-tailed t-test. Three independent biological replicates were used per condition.

<u>DRT10-specific phylogenetic analyses</u>

To identify DRT10 reverse transcriptase homologs, we performed BLASTp searches against the NCBI non-redundant (nr) protein database using representative DRT10 (UG17) RT sequences (WP_123377709.1 and OHD37426.1) as queries (E-value threshold: 0.01, max_target_seqs: 10,000). Retrieved sequences were screened with profile Hidden Markov Models (HMMs) from Mestre et al. (*12*) with hmmscan (HMMER v3.3; E-value < 0.01) (*57*). For each protein sequence, only the top-scoring HMM hit was retained. Sequences matching the DRT10 (UG17) profile (*n* = 3,058) were extracted for downstream analysis.

To reduce computational complexity while maintaining phylogenetic diversity, DRT10 RT sequences were clustered using MMseqs2 (*58*) (easy-linclust mode; minimum sequence identity: 60%, coverage: 80%). After selecting cluster representatives, filtering was applied to remove homologs without retrievable representative loci. The final dataset comprised 492 representative sequences. Multiple sequence alignment was performed using MAFFT (*59*) (v7.505; LINSI option), and a maximum-likelihood phylogenetic tree was constructed using FastTree (*60*) (v2.1.11; -wag -gamma model), before visualization and annotation in iTOL (*61*).

To annotate genetic associations with SLATT, we retrieved flanking genomic regions (±10 kb from RT coding sequences) from NCBI's Identical Protein Groups (IPG) database using the rentrez package in R. Loci were annotated using Prokka (*62*) (v1.14.6; --metagenome mode) with an HMM for SLATT5 that was downloaded from PFAM (PF18160).

ncRNA covariance modeling

Covariance models of ncRNA secondary structure were generated as previously described (*10*). DRT10 RT homologs were identified using the amino acid sequence of the $Eco_1$DRT10, $Eco_2$DRT10, or $Eco_3$DRT10 as the seed query in a BLASTp search on the clusteredNR protein database (max target sequences = 100). Nucleotide sequences 2 kb upstream and downstream of RT genes were retrieved and aligned using MAFFT (v7.505). The resulting alignment was trimmed at the 5′ and 3′ ends to the precise boundaries of the ncRNA as determined by RIP-seq. The resulting sequences were clustered at 99.9% sequence identity to remove duplicates using CD-HIT (*63*) v4.8.1 and realigned using mLocARNA (*64*) v2.0.1 with default parameters. The resulting alignment was used to build and calibrate a covariance model (CM) using the Infernal suite (*65*) v1.1.5. Deduplicated 10 kb genomic contexts for an expanded set of DRT10 RT homologs (see section above; unclustered loci were deduplicated, resulting in 1309 total sequences) were interrogated by the cmsearch function of Infernal to refine the model with additional ncRNAs. The final CM was evaluated and visualized using R-scape (*66*) at an E-value threshold of 0.05 in "Improve given structure" mode.

ncRNA identification and A–B–A' analyses

ncRNA sequences were aligned to clade-specific CMs using cmalign (Infernal v1.1.5) to extract consensus secondary structure annotations. Template loop regions were then identified and extracted via manual inspection of structural annotations and ncRNA alignments. A-B-A' palindromic motifs were identified using a custom R script that searched ungapped sequences for repeated flanking sequences (A and A': 3–5 nt) separated by variable linkers (B: 3–7 nt). Although certain DRT10 homologs exhibit shorter A and A′ regions (as evidenced by $Eco_1$DRT10 and $Eco_2$DRT10), this analysis used a lower length boundary of 3 in order to reduce false positive hits. Motif positions were mapped back to alignment coordinates and collapsed to remove redundant hits with identical midpoint positions within each sequence. Motif distributions were visualized with the ggplot2 package in R. For *She*DRT10 and *Pda*DRT10, A–B–A′ sequences were identified by manual inspection.

RT-SLATT tree of co-evolution

A curated subset of DRT10 RT sequences were clustered with CD-HIT (v4.8.1; 80% amino acid identity), and their associated SLATT sequences were extracted. RT and SLATT sequences were then individually aligned with MAFFT (v7.520; LINSI option), and phylogenetic trees were constructed using FastTree (v2.1.11; -wag -gamma model). Evolutionary similarity was visualized using the cophylo function of the phytools package in R.

Conservation of serine in the DRT10 RT C-terminus

The seven C-terminal residues of RT sequences included in the DRT10-specific tree (described above) were extracted and aligned with the DECIPHER package in R. Positions with less than 50% occupancy were trimmed from the alignment, prior to plotting with the ggplot2 R package and construction of a WebLogo (*67*). The accompanying structural prediction of $Eco_3$DRT10 was obtained with AlphaFold 3 (*68*) and rendered in ChimeraX (*69*).

RT phylogenetic tree analyses

To investigate the phylogenetic relationships between major RT families across all domains of life, we curated a representative collection of all known RTs from the AlphaFold Database. We first collected HMM profiles of RT domains from PFAM (*70*), NCBI CDD (*71*), and TIGRFAM (*72*) databases. In tandem, custom HMM profiles for specific prokaryotic RT subfamilies were compiled from previous phylogenetic analyses (*12*, *73*, *74*). Next, we searched a FASTA-formatted version of the AlphaFold Database (v4) (*75*) using hmmsearch from the HMMER suite (v3.3.2). Hits were filtered to remove any sequences with a domain length below 100 amino acids, E-value above 1e-10, or HMM coverage below 70%. Given that the evolutionary relatedness of RNA-dependent RNA polymerases and maturase K proteins to RTs could result in off-target hits to our HMM profiles (*76*, *77*), we conservatively excluded proteins for which the best hit was an RNA-dependent RNA polymerase (HMM profile PF00978 or any of the NeoRdRP 2.1 HMM profiles (*78*)) or to maturase K (HMM profile CHL00002). Additionally, for each representative target sequence, hits were merged if they derived from the same HMM profile and were fewer than 30 amino acids apart. The RT domain sequences from these 194,634 hits were then extracted by subsetting the hit coordinates with an upstream and downstream buffer of 100 amino acids to ensure domain completeness. Extracted domains were finally clustered at 40% sequence identity using CD-HIT (v4.8.1), yielding 8,533 unique RT domains.

After identifying these RT domains, PDB structures of the corresponding full-length sequences were downloaded from the AlphaFold Database (v4), and RT domains were re-extracted from the PDB structures using the previously determined RT domain coordinates. These domains were then aligned with Reseek (v2.5) (*79*) and Muscle-3D (v5.3) (*80*) using the -super7 algorithm. The resulting structure-based multiple sequence alignment was trimmed to retain only RT motifs 0–7, and positions with >50% gaps were discarded using trimAl (v1.5.rev0) (*81*). To refine the alignment, we removed members with gaps in any of the four conserved positions of the catalytic motif (in RT motif 6), with fewer than 100 amino acids aligned, or with partial RT domains (where the alignment started after RT motif 1 or ended before RT motif 7). Finally, a maximum-likelihood phylogenetic tree was constructed from the resulting alignment of 6,734 RT domain sequences using VeryFastTree (v4.0.5) (*82*) (-wag -pseudo -gamma). The RT sequences used in this phylogenetic analysis are provided in **Table S4**. The multiple sequence alignment of the RT domains, the phylogenetic tree, and the tree annotations are provided as **Data S1-S3**. See **Supplementary Text** for a detailed explanation of methodological considerations.

RT structure-based neighbor-joining tree analysis

To supplement the maximum-likelihood phylogenetic tree, a neighbor-joining tree from pairwise structural alignments was built, as follows. All-vs-all comparisons were first performed using Foldseek (*83*) (commit version 1d17b43) (--exhaustive-search 1 --alignment-type 1 --exact-tmscore 1 --max-seqs 10000) on the full-length PDB structures retrieved from the phylogenetic tree analysis described above. Then, the inverse of the normalized structural bit score was used to build a distance matrix that was subsequently fed into DecentTree (v1.0.0) (*84*) to construct a neighbor-joining tree with the NJ-R algorithm. The neighbor-joining tree is provided as **Data S4**.

RT structural perplexity analysis

To determine the closest structural neighbors to human TERT (hTERT), we employed an information-theoretic measure of structural distance between two proteins called structural perplexity (*32*). First, we manually compiled a reference catalog of major RT families across all domains of life, including $Eco_1$DRT10, $Eco_2$DRT10, and $Eco_3$DRT10 (**Table S5**). When available, structural predictions from the AlphaFold Database (v4) were retrieved using AlphaFetcher. For RTs without structures available in the AlphaFold Database, structures were predicted using AlphaFold 3. Then, all protein structures were aligned against hTERT (O14746) using MMLigner (v1.0.2) (*85*), and the structural perplexity was computed as $2^{-\text{Compression}}$ per the Plexy algorithm (*32*) (**Table S6**). Finally, all structures that yielded negative compression values were excluded from the analysis, and only those with structural perplexity values of less than 1e-10 were visualized.

Mammalian cell culture

HEK293T cells were purchased from the American Type Culture Collection (ATCC CRL-3216). Cells were cultured at 37 °C and 5% $CO_2$ in DMEM with 10% FBS and 100 U $mL^{-1}$ of penicillin and streptomycin (Thermo Fisher Scientific) and were routinely tested for mycoplasma contamination.

RIP-seq and cDIP-seq in human cells

HEK293T cells were seeded in 10 cm dishes with 2 × $10^6$ cells per dish. After 24 hours, cells were co-transfected with separate plasmids encoding the *Eco*$_3$DRT10 3xFLAG-tagged RT and the ncRNA using Lipofectamine 2000 (Thermo Fisher Scientific), per the manufacturer's instructions. Cells were harvested 72 hours after transfection by aspirating culture media, detaching cells with trypsin, and collecting cell pellets by centrifugation at 2,000 × *g* for 5 min at 4 °C. Pellets were washed with 1 mL of PBS and centrifuged again at 2,000 × *g* for 2 min at 4 °C. After supernatants were removed, pellets were flash-frozen in liquid nitrogen and temporarily stored at –80 °C.

Immunoprecipitation was performed as described above for *E. coli* cells, with minor modifications. Antibody–bead complexes were prepared for immunoprecipitation as above, by washing Dynabeads Protein G (60 µL per sample) three times with 1 mL Human RIP Dilution Buffer (50 mM Tris pH 7.5, 150 mM NaCl, 1 mM $MgCl_2$, 0.05% NP-40), resuspending in 1 mL Human RIP Dilution Buffer, and combining with anti-FLAG antibody (20 µL per sample) before rotating for > 3 hours at 4 °C. Antibody–bead complexes were then washed twice and resuspended in Human RIP Dilution Buffer to a final volume of 60 µL per sample. Flash-frozen cell pellets were thawed on ice and resuspended in 100 µL Human RIP Lysis Buffer (10 mM HEPES-KOH pH 7.0 at 25 °C, 100 mM KCl, 5 mM $MgCl_2$, 0.5% NP-40) supplemented with 1 mM DTT, 1× Complete Protease Inhibitor Cocktail (Roche), and 0.1 U $\mu L^{-1}$ SUPERase•In RNase Inhibitor (Thermo Fisher Scientific). After 5 minutes on ice, lysates were frozen at –80 °C. Frozen lysates were thawed on ice and centrifuged at 16,000 × *g* for 10 min at 4 °C to pellet cell debris. Supernatants were transferred to new tubes and diluted 1:10 in Human RIP Dilution Buffer supplemented with 1 mM DTT, 1× Complete Protease Inhibitor Cocktail, and 0.1 U $\mu L^{-1}$ SUPERase•In RNase Inhibitor. A 10 µL aliquot of each sample was set aside as an "input" control and stored at –80 °C. The remainder of each sample was combined with 60 µL of antibody-bead complex and rotated overnight at 4 °C. The next day, each sample was washed 3× with 1 mL ice-cold Human RIP Dilution Buffer. During the final wash, each sample was split into two 500 µL volumes for downstream RIP or cDIP processing. RIP elutions were performed as described above for *E. coli* samples. For cDIP elutions, beads were resuspended in 80 µL IP Wash Buffer and treated with 5 µg RNase A (Thermo Fisher Scientific) at 37 °C for 30 min. SDS was then added to each sample to a final concentration of 1% before treating with 25 µg Proteinase K (Thermo Fisher Scientific) at 55 °C for 30 min. Beads were pelleted using a magnetic rack, and supernatants containing eluted DNA were transferred to new tubes. DNA was isolated using the Monarch Spin PCR and DNA Cleanup kit (NEB), following the Oligonucleotide Cleanup protocol. RIP-seq HTS libraries were prepared as described above for *E. coli* cells, and cDIP-seq HTS libraries were prepared as described for miniprep-seq samples. Libraries were sequenced on an Element AVITI in single-end mode with 150 cycles.

RIP-seq samples were analyzed as described above for experiments from *E. coli* cells, mapping reads to a combined reference file for the RT and ncRNA expression plasmids. cDIP-seq reads were trimmed and filtered as described above for miniprep-seq. Datasets were randomly downsampled to 100,000 reads per sample using seqtk, and the resulting files were analyzed to generate motif position graphs. Motif counts were normalized to counts per million reads by multiplying by a scaling factor of 10.

### SUPPLEMENTARY TEXT

Methodological considerations for phylogenetic reconstruction

Reverse transcriptases (RTs) exhibit remarkable sequence diversity. This is unsurprising, given that their frequent mobilization, their presence across all domains of life, and their likely ancient origin (*86*, *87*).

As most methods for phylogenetic reconstruction are based on multiple sequence alignments (MSAs), such sequence divergence makes accurate phylogenetic inference particularly challenging. Firstly, substitutional saturation can obscure or underestimate true divergence, compressing long branches and promoting a long-branch attraction artifact. Secondly, such divergence also leaves few confidently homologous positions, so stochastic error and alignment uncertainty can introduce spurious synapomorphies that bias branch lengths and topology.

To improve the quality of the MSA underlying our RT phylogeny, we developed a methodology to minimize errors derived from aligning these highly divergent sequences. First, we combined a comprehensive RT domain search with stringent filtering criteria to ensure domain completeness. Second, we leveraged structure-based MSAs that have previously been used to reconstruct the evolutionary history of other ancient enzymes with high sequence divergence, such as B-family DNA polymerases (*88*) and RNA-dependent RNA polymerases (*89*). Third, we conservatively trimmed highly gapped positions and removed spurious sequences to maximize reliable homology while retaining informative variability. Below, we provide a detailed explanation of these decisions.

Detailed methodology for RT phylogenetic analysis

*Comprehensive RT domain search that avoided truncated domains*

To assemble a comprehensive dataset of RTs across all domains of life, we used hmmsearch from the HMMER suite to screen a FASTA-formatted version of the AlphaFold Database (*75*) with Hidden Markov Models (HMMs) of RT domains from the public PFAM (*70*), NCBI CDD (*71*), and TIGRFAM (*72*) databases, as well as custom HMM profiles for specific prokaryotic RT subfamilies reported in recent phylogenies (*12*, *73*, *74*).

To maintain a high level of confidence in the retrieved hmmsearch hits, we applied several stringent filtering steps. First, we ensured high-quality hits by removing any with a domain length below 100 amino acids, E-value above 1e-10, or HMM coverage below 70%. Second, we conservatively removed proteins for which the best hit was either an RNA-dependent RNA polymerase (HMM profile PF00978 or any of the NeoRdRP 2.1 HMM profiles (*78*)) or a maturase K protein (HMM profile CHL00002), because these enzymes may yield significant hits in RT HMM profiles due to their evolutionary relatedness (*76*, *77*).

To compile a dataset of RT domains that were as complete as possible, we merged overlapping or adjacent RT hits originating from the same HMM profile and separated by fewer than 30 amino acids for each representative. Such small gaps typically reflect alignment discontinuities due to insertions rather than distinct domains. We then extracted the sequences delineated by the hmmsearch hit coordinates while adding an upstream and downstream buffer of 100 amino acids to ensure complete domain capture, since hmmsearch hit sequences can truncate prematurely. This yielded a large collection of putative complete RT domains, which were clustered permissively at a 40% sequence identity threshold using CD-HIT to create a non-redundant set spanning the diversity of RTs. The resulting 8,533 clusters formed the basis of our unique RT domain dataset.

*Structure-based MSA of the RT domain*

To align these RT domains and build a high-quality MSA for our phylogenetic analysis, we circumvented the high sequence divergence of RTs by avoiding sequence-only based alignment methods and opted for a structure-based MSA instead, which more accurately aligns proteins with low sequence identity and remote homology (*76*, *77*, *90*). We first retrieved the predicted structures of full-length RT sequences corresponding to each RT domain in our dataset from the AlphaFold Database. We then trimmed the coordinates to the same domain boundaries used in the primary sequence-level extraction. This provided a one-to-one mapping between each RT domain sequence and its predicted structure. Finally, we generated structure-based multiple sequence alignments (MSAs) using Reseek (*79*) and Muscle-3D with the -super7 algorithm (*80*). We used Muscle-3D because it scales well to thousands of proteins, compared to other recently published structure alignment methods.

*Processing the MSA to ensure high-quality alignment of complete RT domains*

To ensure that the resulting MSA retained only the most reliably aligned positions, it was first trimmed to include only the canonical RT motifs 0–7. Any column with more than 50% gaps was then pruned using trimAl (*81*). To further eliminate spurious or incomplete entries, we also stringently removed any MSA members with gaps at any of the four conserved catalytic residues in RT motif 6, an alignment length shorter than 100 amino acids, or a partial core RT domain that begins after RT motif 1 or ends before RT motif 7.

Altogether, these criteria ensured that only high-quality, full-length RT domains with catalytic motifs were included in the final MSA of 6,734 members, allowing us to reduce artifacts arising from alignment errors and increase the robustness of the downstream maximum-likelihood phylogenetic tree, which we generated using VeryFastTree (*82*).

<u>Choice of tree-building algorithm</u>

Pythia (*91*), a tool that quantifies the difficulty of maximum-likelihood-based tree inferences for a given MSA, classified our alignment as “difficult” (difficulty = ~0.7). This scoring predicts that starting trees (or seeds) can produce multiple near-equal-likelihood final topologies, and that convergence can be slow and ambiguous. At this level of difficulty, different maximum-likelihood phylogenetic tools such as FastTree, IQ-TREE, and RAxML may yield differing topologies at similar likelihoods, though IQ-TREE and RAxML are often regarded as more accurate at lower dataset difficulties (*92*).

Given this parity in tree-building accuracy, we chose to employ VeryFastTree, a highly optimized implementation of FastTree specifically designed to align massive numbers of sequences (*82*), as the large size of our dataset was causing slow convergence of IQ-TREE and RAxML trees. Additionally, the choice of a FastTree-based method (with WAG+Γ and pseudocounts) was well supported by literature precedent from previous large-scale analyses of prokaryotic RTs (*12*, *73*, *74*, *93–95*).

<u>Orthogonal support</u>

To complement our phylogenetic analysis, we further employed two parallel structure-based computational analyses.

*Structure-based neighbor-joining tree*

We first built a structure-based tree by running all-vs-all Foldseek comparisons on the full-length AlphaFold models, converting normalized bit scores into distances, and reconstructing the tree using DecentTree (NJ-R). Because full-length comparisons incorporate accessory domains (e.g., RNase H, fingers/thumb extensions, lineage-specific inserts, etc.) in addition to the core RT domain, this analysis assesses overall architectural similarity rather than the core palm domain alone. Despite potential noise from flexible/disordered regions, the structure-based tree recovered broad partitions precisely where the sequence signal is weakest (i.e., resolved canonical group II intron RTs, retron-like RTs, DGR-associated RTs, and LTR/non-LTR lineages). This structure-distance clustering analysis provides an orthogonal line of evidence to our phylogenetic analysis for the broad assignment of RT groupings. Indeed, agreement between the maximum-likelihood phylogenetic tree and this structure-distance clustering analysis at coarse scales increases confidence that proposed groupings are not solely due to alignment artifacts or model misspecification.

*Structural perplexity analysis*

We further leveraged an information-theoretic measure of structural distance between two proteins called structural perplexity to compare human TERT (hTERT) against a curated catalog of representative RTs across domains of life. Performing this analysis using MMLigner (*85*) and the Plexy algorithm (*32*), where the structural perplexity was computed as $2^{-\text{Compression}}$, once again recapitulated the finding of structural similarity between TERT and bacterial RTs, including Class 2 UG proteins.

In summary, we present a phylogenetic reconstruction of RT evolution that is supported by a structure-based MSA, structure-based distances from comparing full-length structural models, and detailed mechanistic evidence. Despite the challenges laid out above in the phylogenetic analysis of highly diverged proteins, the convergence of multiple lines of evidence in our study reinforces our conclusion that TERT shares a close evolutionary relationship with UG RTs, and that RNA-templated cDNA repeat addition most likely arose in a bacterial ancestor of telomerase.

### SUPPLEMENTARY FIGURES

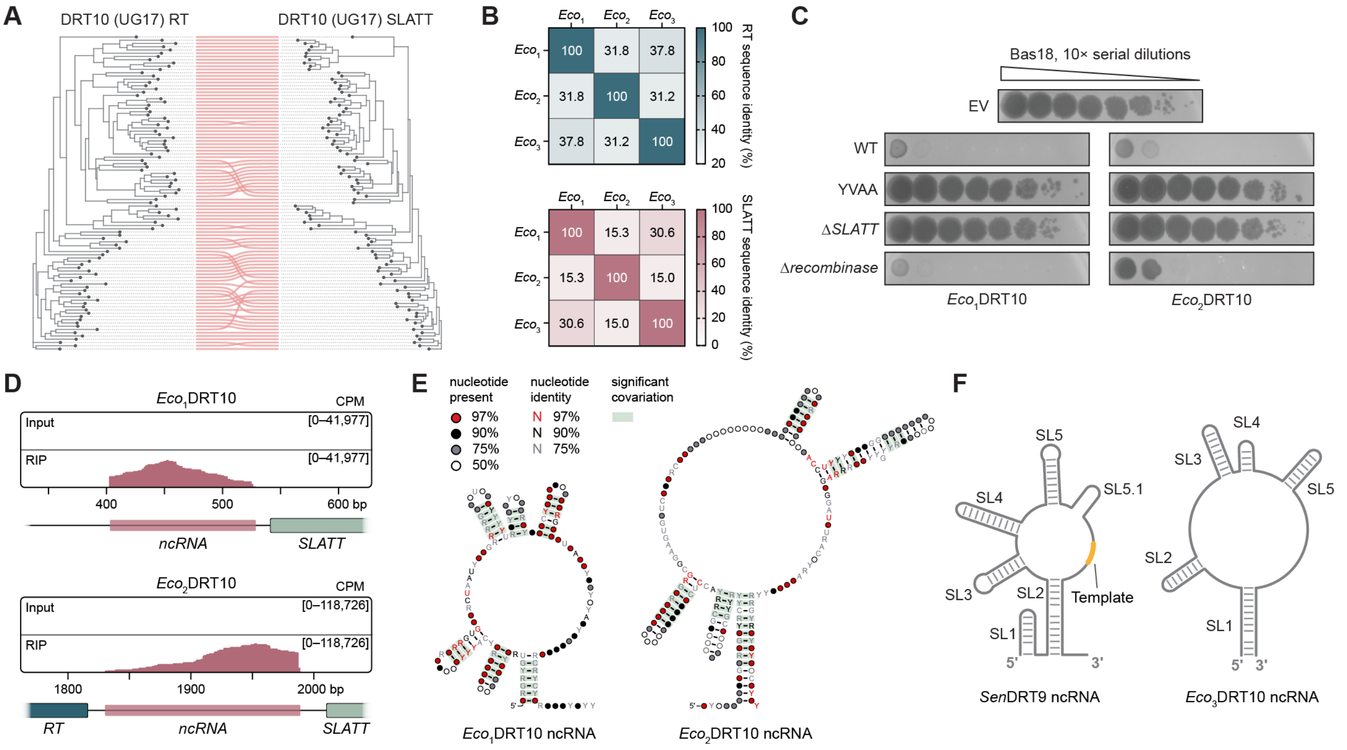


**Figure S1. RNA and protein components of DRT10 phage defense systems. (A)** Comparison of phylogenetic trees for select DRT10 (UG17) RT and SLATT homologs, revealing strong genetic linkage and co-evolution. **(B)** Sequence identity matrices for RT proteins (top) and SLATT proteins (bottom) from DRT10 systems selected for experimental study. **(C)** Plaque assays for *Eco*$_1$DRT10 and *Eco*$_2$DRT10 systems demonstrating robust defense against Bas18 phage, which is lost with mutation of the RT catalytic residues (YVDD>YVAA) or deletion of the SLATT gene (Δ*SLATT*). Deletion of a tyrosine recombinase gene encoded nearby (Δ*recombinase*) has no effect on defense activity. **(D)** RIP-seq coverage tracks for *Eco*$_1$DRT10 (top) and *Eco*$_2$DRT10 (bottom) RTs compared to non-immunoprecipitated ("Input") controls. Schematics of the genetic loci from the corresponding expression plasmids are shown below; data are normalized for sequencing depth and plotted as counts per million reads (CPM). **(E)** Covariance models of the *Eco*$_1$DRT10 (left) and *Eco*$_2$DRT10 (right) ncRNA secondary structures from analyses of 283 and 159 homologous systems, respectively. **(F)** Comparison of the experimentally determined ncRNA secondary structure for *Sen*DRT9 and the predicted ncRNA secondary structure for *Eco*$_3$DRT10.

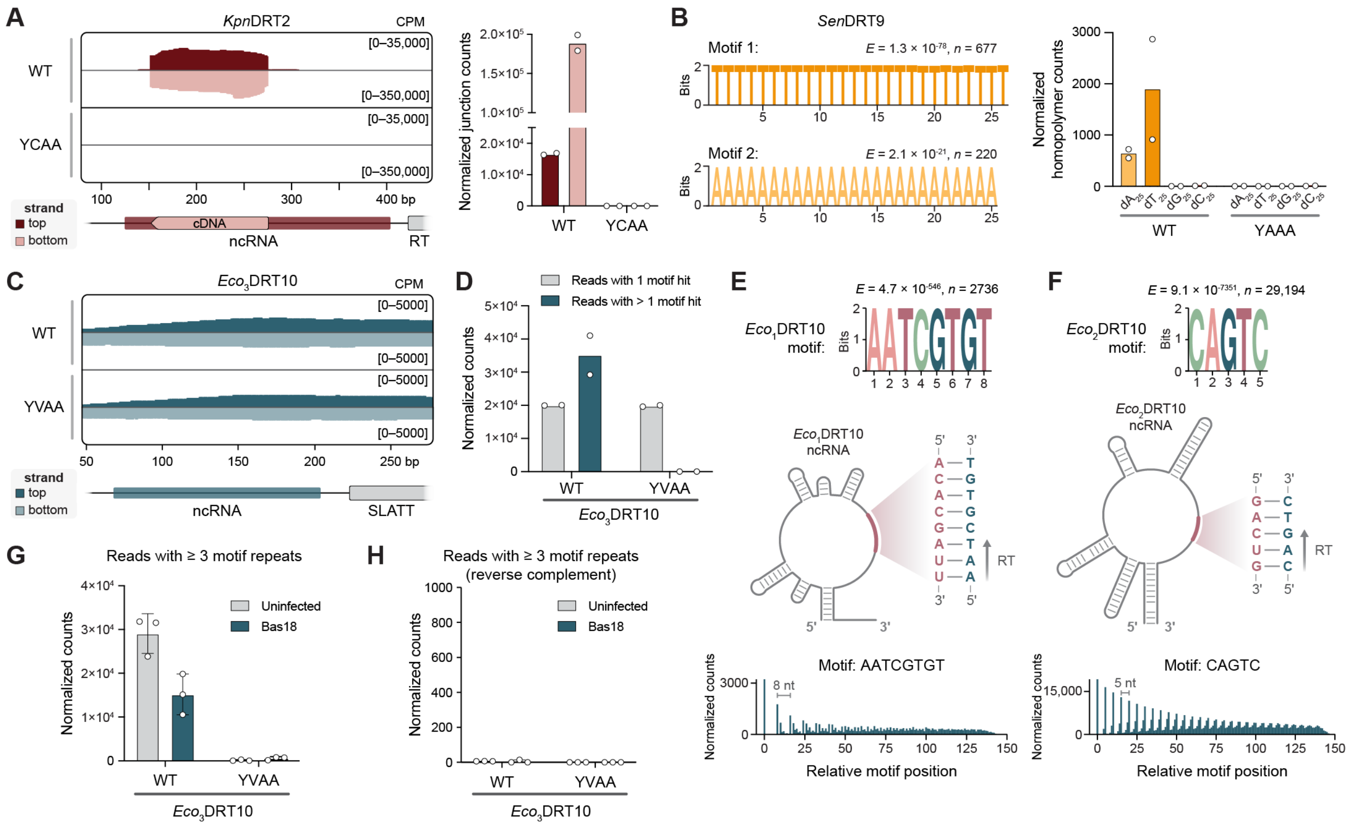


**Figure S2. DRT10 synthesizes tandem-repeat cDNA using an RNA template. (A)** *Left*: Coverage tracks for miniprep-seq of *E. coli* cells expressing WT or catalytically inactive (YCDD>YCAA) *Kpn*DRT2 during T5 phage infection, shown over the plasmid-encoded ncRNA locus. Top-strand and bottom-strand alignments are shown above and below the axis, respectively; data are normalized for sequencing depth as counts per million reads (CPM). *Right*: Quantification of top-strand and bottom-strand miniprep-seq reads spanning the cDNA repeat junction. Data are shown for *n* = 2 independent biological replicates. **(B)** *Left*: MEME motif discovery results for unmapped miniprep-seq reads from *E. coli* cells expressing *Sen*DRT9 during T5 phage infection. Motifs significantly enriched in WT samples relative to catalytically inactive (YADD>YAAA) control samples are shown. *E* represents the *E*-value significance; *n* represents the number of contributing sites. *Right*: Quantification of homopolymer-containing reads from WT and YAAA miniprep-seq datasets. Data are shown for *n* = 2 independent biological replicates and are normalized as in **A**. **(C)** Coverage tracks for miniprep-seq of *E. coli* cells expressing WT or catalytically inactive (YVAA) *Eco*$_3$DRT10 in the absence of phage. Top-strand and bottom-strand alignments are shown over the plasmid-encoded ncRNA locus and are normalized as in **A**. **(D)** Bar graph quantifying the number of *Eco*$_3$DRT10 miniprep-seq reads containing exactly 1 or > 1 hit to the motif identified in **Fig. 2B**. Reads containing only 1 motif hit likely derive from the plasmid. **(E)** *Top*: MEME motif discovery results for unmapped miniprep-seq reads from cells expressing *Eco*$_1$DRT10, showing the top enriched motif for WT samples relative to a YVAA control. *E* represents the *E*-value significance; *n* represents the number of contributing sites. *Middle*: Schematic of the *Eco*$_1$DRT10 ncRNA secondary structure and an inset highlighting the likely template region, which matches the DNA motif identified above as its reverse complement. *Bottom*: Motif position graph for miniprep-seq of cells expressing *Eco*$_1$DRT10, showing 8-nt periodicity of the motif identified above. **(F)** MEME motif discovery, ncRNA template and motif schematic, and motif position graph for *Eco*$_2$DRT10, shown as in **E. (G)** Bar graph quantifying the number of *Eco*$_3$DRT10 miniprep-seq reads containing ≥ 3 consecutive repeats of the motif identified in **Fig. 2B**, for WT and YVAA samples in the absence or presence of Bas18 phage. Data are mean ± SD for *n* = 3 independent biological replicates. **(H)** Bar graph quantifying the number of *Eco*$_3$DRT10 miniprep-seq reads containing ≥ 3 consecutive repeats of the reverse complement of the 9-nt motif, for WT and YVAA samples in the absence or presence of Bas18 phage. Data are mean ± SD for *n* = 3 independent biological replicates. For **D–H**, data are normalized as counts per million plasmid-mapping reads.

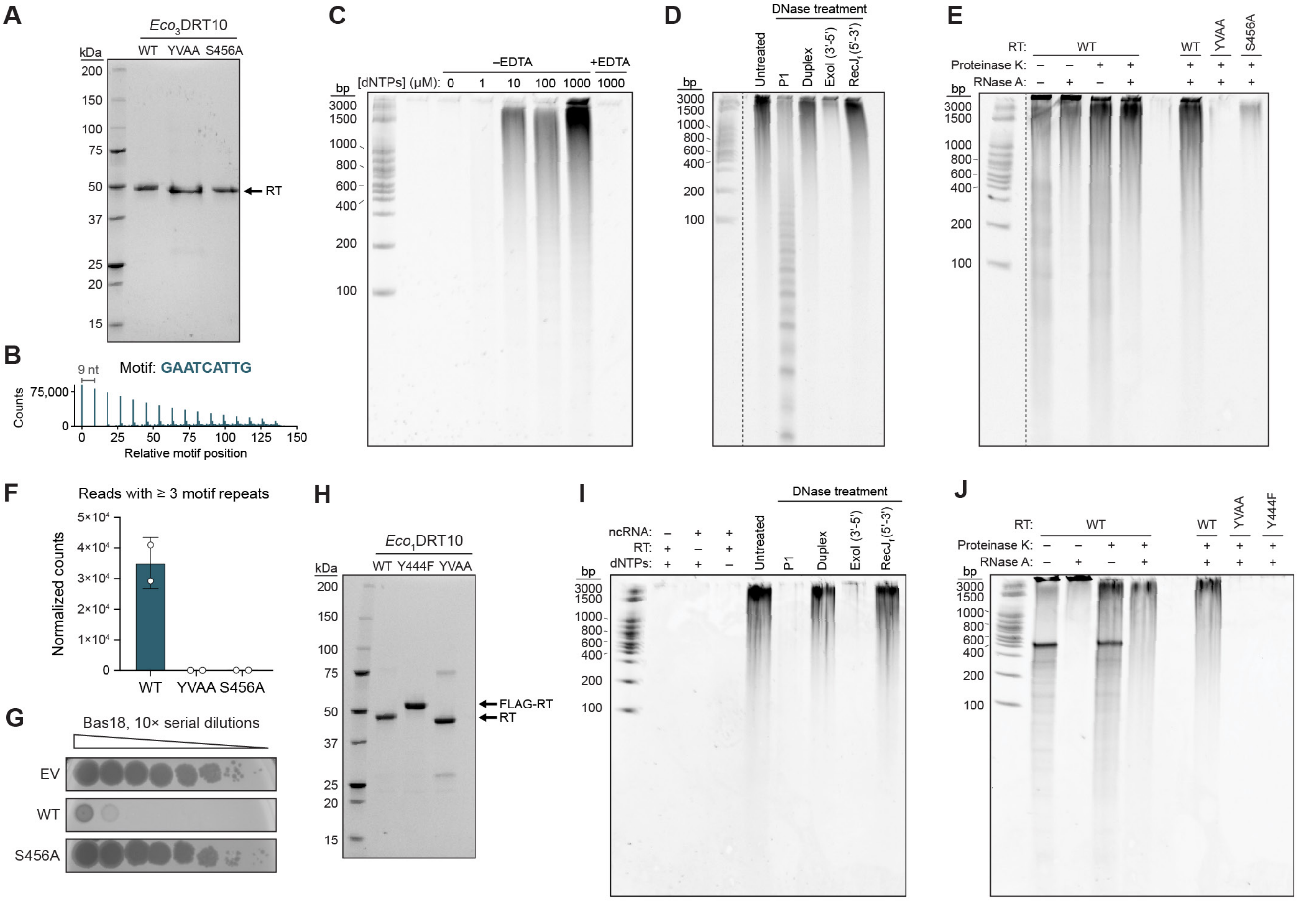


**Figure S3. Biochemical requirements for protein-primed DNA synthesis by DRT10. (A)** Denaturing 10% SDS-PAGE analysis of purified *Eco*$_3$DRT10 RT and the indicated mutants. Sizes from a protein ladder are shown. **(B)** Motif position graph from high-throughput sequencing of *in vitro* DNA synthesis products from reactions containing purified *Eco*$_3$DRT10 RT, ncRNA, and dNTPs. **(C)** Denaturing 15% urea-PAGE analysis of *Eco*$_3$DRT10 RT activity assays investigating the dNTP- and metal ion-dependence of cDNA synthesis. All reactions contained 2.5 µM RT–ncRNA complexes and were incubated at 37 °C for 60 min with the indicated dNTP concentration; EDTA, when present, was added to a final concentration of 10 mM prior to dNTP addition. Reactions were terminated by boiling and treated with proteinase and RNase prior to electrophoretic analysis. Sizes from a dsDNA ladder are shown. **(D)** Denaturing 15% urea-PAGE analysis of *Eco*$_3$DRT10 RT activity assays evaluating the sensitivity of RT reaction products to different nuclease treatments. All reactions contained 2.5 µM RT–ncRNA complexes and were incubated with 100 µM dNTPs at 37 °C for 60 minutes prior to enzymatic treatment. Nuclease P1 is a ssDNA-specific endonuclease; Duplex DNase is a dsDNA-specific endonuclease; ExoI is a 3′-5′ ssDNA exonuclease; RecJ$_f$ is a 5′-3′ ssDNA exonuclease. Sizes from a dsDNA ladder are shown. **(E)** Denaturing 15% urea-PAGE analysis of *Eco*$_3$DRT10 RT activity assays with WT RT and the indicated variants, treated with proteinase and RNase as indicated. Reactions were performed as in **D**; sizes from a dsDNA ladder are shown. **(F)** Bar graph quantifying the number of miniprep-seq reads containing ≥ 3 repeats of the DNA motif for *Eco*$_3$DRT10 WT RT, compared to RT variants with mutations in the active site (YVAA) or putative priming residue (S456A). Data are normalized for sequencing depth as counts per million plasmid-mapping reads, and are shown for *n* = 2 independent biological replicates. **(G)** Plaque assays for *Eco*$_3$DRT10 against Bas18 phage, comparing the WT system and S456A mutant to an empty vector (EV) control. Mutation of the putative priming residue (S456) leads to a complete loss of defense. **(H)** Denaturing 10% SDS-PAGE analysis of purified *Eco*$_1$DRT10 RT, including the WT protein, an active site mutant (YVAA) and a FLAG-tagged priming residue mutant (Y444F). Sizes from a protein ladder are shown. **(I)** Denaturing 15% urea-PAGE analysis of *Eco*$_1$DRT10 RT activity assays evaluating the sensitivity of RT reaction products to different nuclease treatments, performed as in **D**. **(J)** Denaturing 15% urea-PAGE analysis of *Eco*$_1$DRT10 RT activity assays with WT RT and the indicated variants, performed as in **E**. The *Eco*$_1$DRT10 ncRNA is indicated.

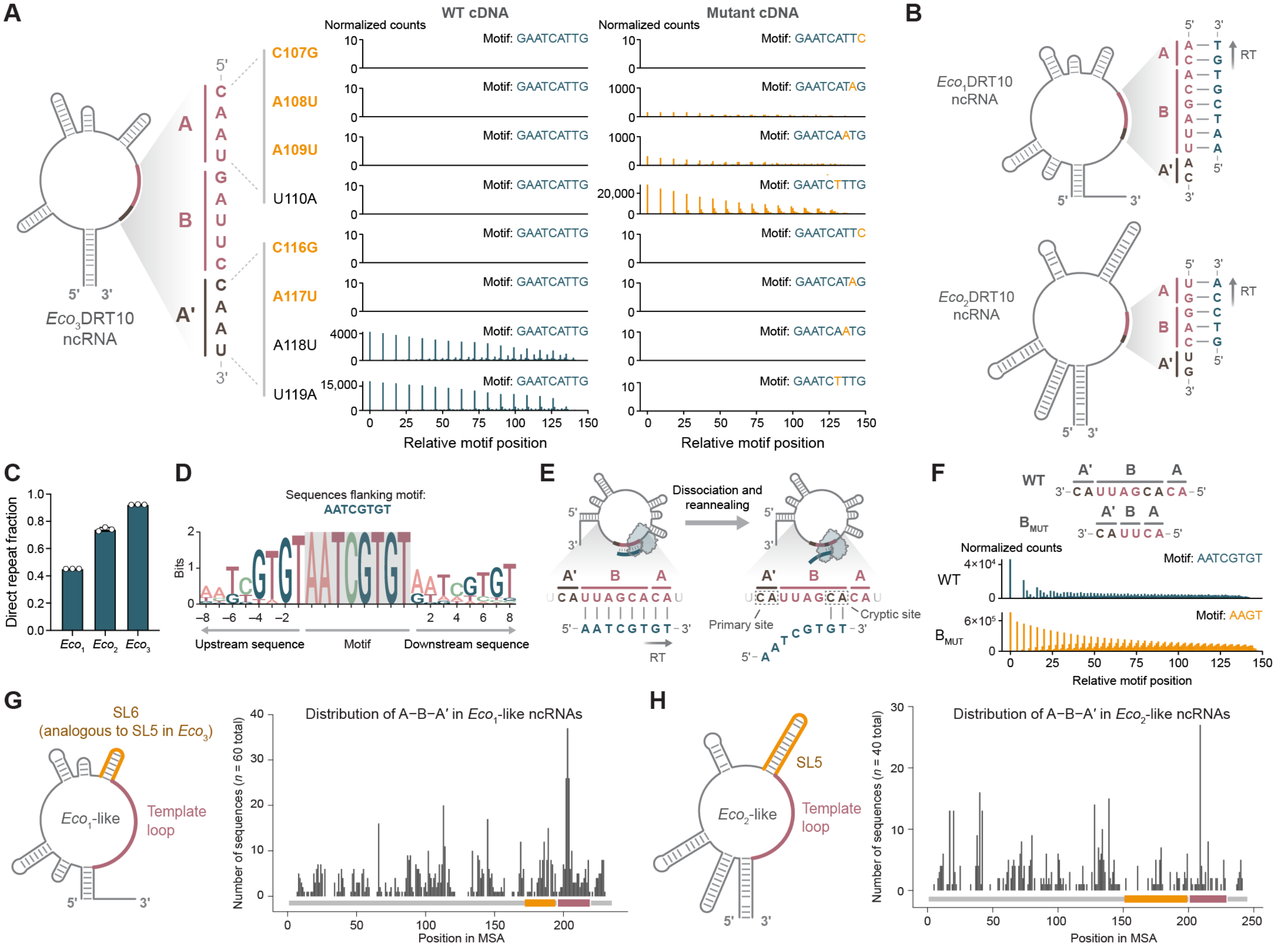


**Figure S4. RNA determinants of DRT10-mediated cDNA repeat addition. (A)** *Left*: Illustration of *Eco*$_3$DRT10 ncRNA with the template region highlighted in red. An inset shows the A–B–A′ repeat pattern within the template region. *Right*: Motif position graphs from miniprep-seq datasets of the indicated *Eco*$_3$DRT10 ncRNA variants; mutations associated with a loss of phage defense are bolded orange. The first column shows the position of WT cDNA motif hits, while the second column shows the position of mutant cDNA motif hits that would be generated by each template mutant. Data are normalized for sequencing depth as counts per million plasmid-mapping reads. **(B)** Illustrations of the *Eco*$_1$DRT10 (top) and *Eco*$_2$DRT10 (bottom) ncRNAs, with insets highlighting the A–B–A′ patterns within their respective template regions. Note that the cDNA repeat sequence templated by A–B for *Eco*$_2$DRT10 is a permutation of the motif identified in **Fig. S2F**. **(C)** Bar graph quantifying the "direct repeat fraction," or the fraction of motif hits in DRT10 miniprep-seq reads that are precisely flanked on either side by motif repeats. *Eco*$_1$DRT10 has the lowest direct repeat fraction of the tested homologs, indicative of inaccurate repeat addition. Data are mean ± SD for $n = 3$ independent biological replicates. **(D)** Sequence logo of motif hits and flanking sequences from *Eco*$_1$DRT10 miniprep-seq datasets, showing the degeneracy of motif-flanking sequences. **(E)** Schematic of incomplete repeat addition by *Eco*$_1$DRT10 due to the presence of a cryptic re-annealing site within B, which leads to the addition of a 2-nt 5′-GT-3′ sequence rather than the full 8-nt motif in subsequent rounds of extension. **(F)** Motif position graphs from *Eco*$_1$DRT10 miniprep-seq datasets with the template sequences schematized at top. The WT template contains the cryptic re-annealing site, leading to incomplete repeat addition and correspondingly weak periodicity in the motif position graph. Mutation of B to remove this cryptic site results in efficient synthesis of a 4-nt cDNA repeat motif with strong periodicity. Data are normalized for sequencing depth as counts per million plasmid-mapping reads. **(G,H)** Distribution of bioinformatically predicted A–B–A′ motifs in *Eco*$_1$-like (**G**) and *Eco*$_2$-like (**H**) ncRNA sequences. Motif midpoints are plotted non-redundantly, visualizing one motif midpoint per ncRNA.

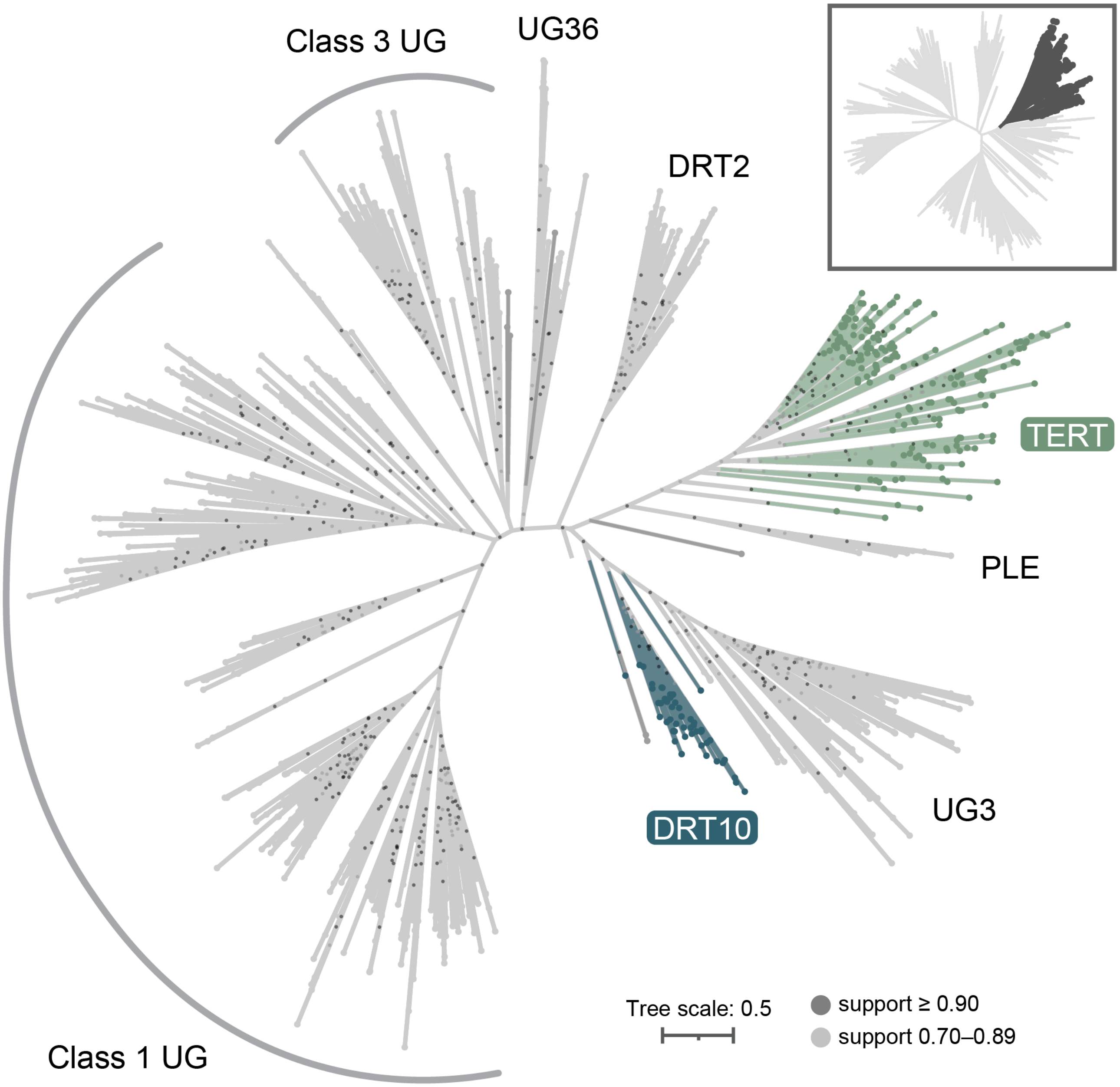


**Figure S5. Phylogenetic relationships between UG and TERT family RTs.** Pruned subtree derived from the larger phylogenetic tree in **Figure 4B,** as shown by the highlighted clade in the inset at top right. RT sequences belonging to the well-supported clade (local support value: 0.987) comprising TERT, PLE, and UG RTs were retained, enabling closer inspection of their evolutionary relationships.

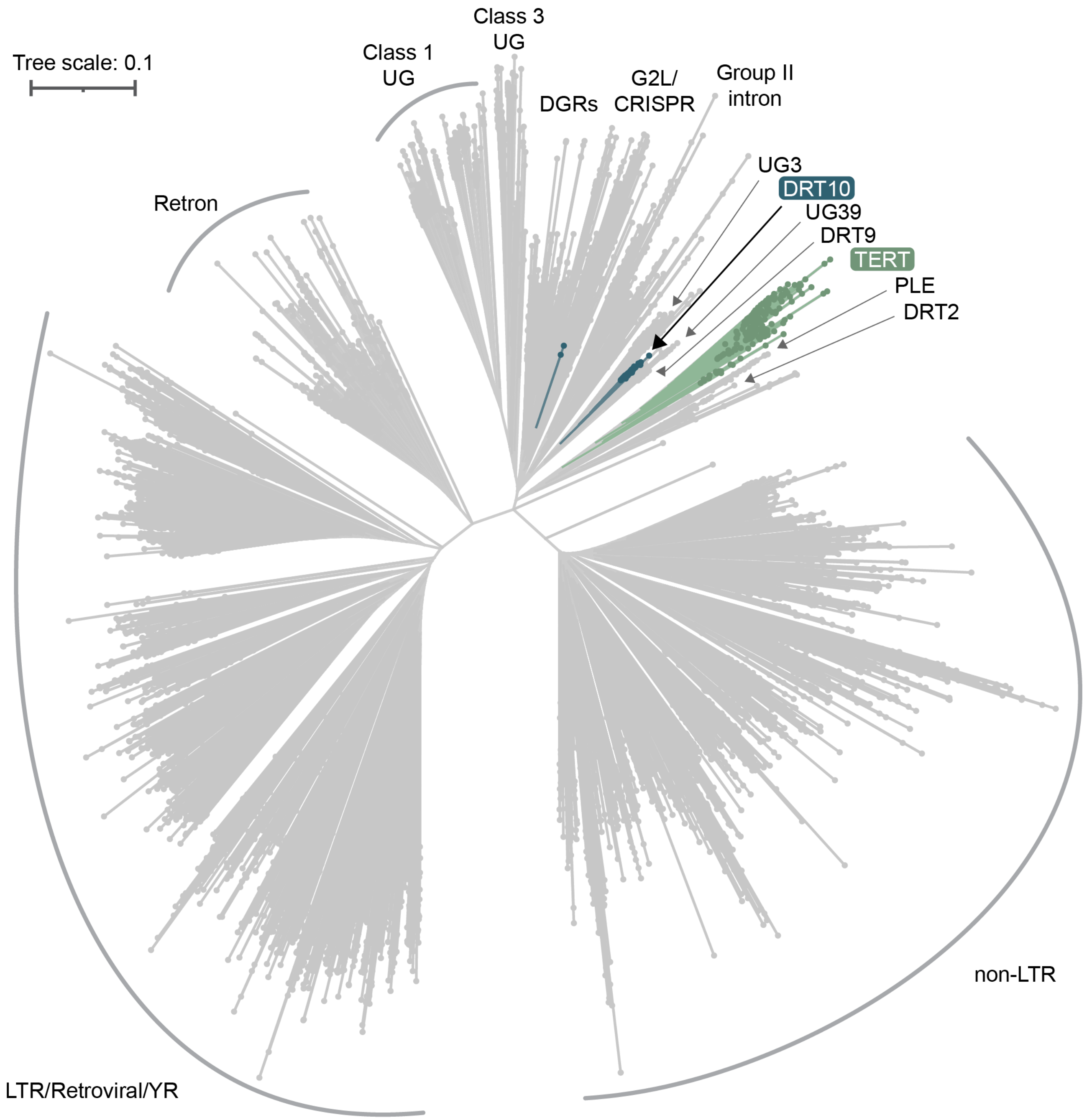


**Figure S6. Structure-based neighbor-joining tree of major RT families across the three domains of life.** Neighbor-joining (NJ) tree based on Foldseek comparisons of structural models for all RTs shown in **Fig. 4B**. Distances in the tree represent the inverse of the structural bit score rendered by Foldseek. The NJ tree was built with DecentTree using the NJ-R algorithm.

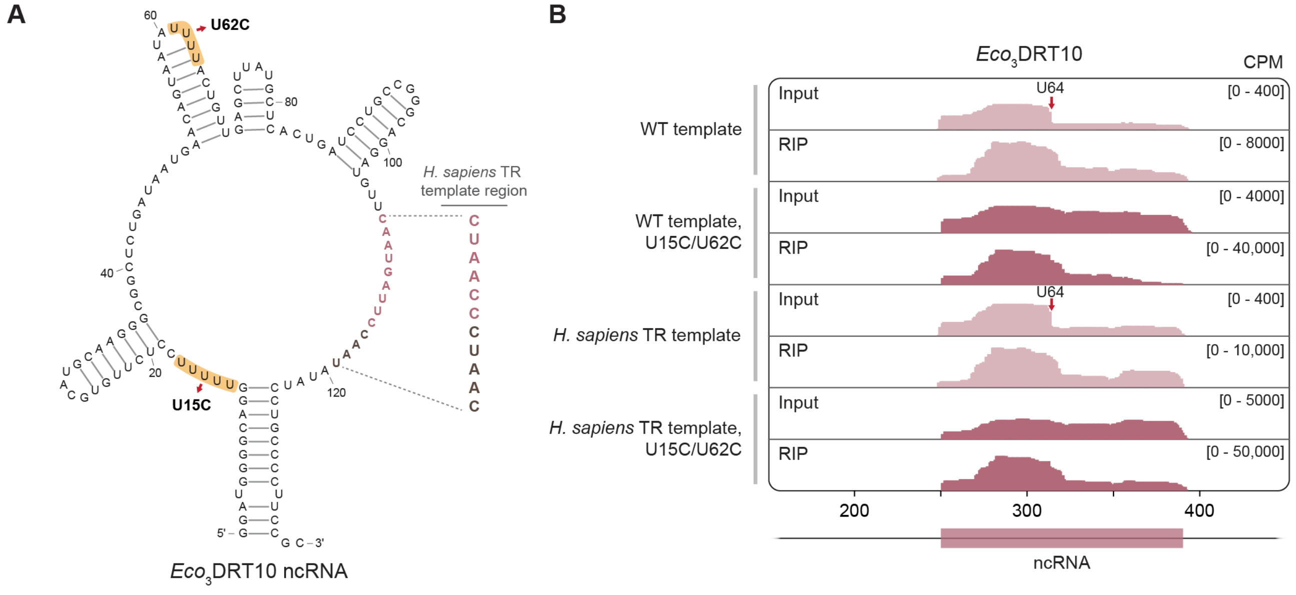


**Figure S7. Optimization of *Eco*$_3$DRT10 ncRNA expression in human cells. (A)** Schematic of the *Eco*$_3$DRT10 ncRNA secondary structure, with poly-U tracts highlighted in yellow. Mutations designed to improve ncRNA expression in human cells by RNA polymerase III, which terminates at poly-U tracts, are indicated. The template region is bolded in red (A–B) and brown (A′), and the *H. sapiens* TR template is shown at right. **(B)** Coverage tracks from RIP-seq datasets of FLAG-tagged *Eco*$_3$DRT10 RT expressed in HEK293T cells together with the indicated ncRNA variants, shown over the plasmid-encoded ncRNA locus. A drop-off in coverage is seen after U64 for the unmodified ncRNA, indicative of partial transcription termination triggered by the poly-U tract. Disruption of the poly-U tracts with U15C/U62C mutations results in increased expression of the full-length ncRNA and stronger association with the RT. Data are normalized for sequencing depth as counts per million reads (CPM).

#### SUPPLEMENTARY TABLES

**Table S1.** DRT10 sequences in Fig. 3H phylogenetic tree.

**Table S2.** Bioinformatically predicted DRT10 ncRNA sequences.

**Table S3.** Bioinformatically predicted A–B–A′ patterns in DRT10 ncRNA sequences.

**Table S4.** RT sequences in phylogenetic and structure-based neighbor-joining trees.

**Table S5.** RT sequences in structural perplexity analysis.

**Table S6.** Results of structural perplexity analysis against human TERT.

**Table S7.** Strains used in this study.

**Table S8.** Descriptions and sequences of plasmids used in this study.

**Table S9.** Experimentally studied DRT10 homologs.

#### SUPPLEMENTARY DATA

**Data S1.** Multiple sequence alignment of RT domains for the Fig. 4B phylogenetic tree.

**Data S2.** Annotation file for Data S3 and Data S4 tree files.

**Data S3.** Phylogenetic tree in Fig. 4B.

**Data S4.** Structure-based neighbor-joining tree in fig. S6.
